# SArKS: *de novo* discovery of gene expression regulatory motifs and domains by suffix array kernel smoothing

**DOI:** 10.1101/133934

**Authors:** Dennis Wylie, Hans A. Hofmann, Boris V. Zemelman

## Abstract

**Motivation:** We set out to develop an algorithm that can mine differential gene expression data to identify candidate cell type-specific DNA regulatory sequences. Differential expression is usually quantified as a continuous score—fold-change, test-statistic, *p*-value—comparing biological classes. Unlike existing approaches, our *de novo* strategy, termed SArKS, applies nonparametric kernel smoothing to uncover promoter motifs that correlate with elevated differential expression scores. SArKS detects motifs by smoothing sequence scores over sequence similarity. A second round of smoothing over spatial proximity reveals multi-motif domains (MMDs). Discovered motifs can then be merged or extended based on adjacency within MMDs. False positive rates are estimated and controlled by permutation testing.

**Results:** We applied SArKS to published gene expression data representing distinct neocortical neuron classes in *M. musculus* and interneuron developmental states in *H. sapiens*. When benchmarked against several existing algorithms for correlative motif discovery using a cross-validation procedure, SArKS identified larger motif sets that formed the basis for regression models with higher correlative power.

**Availability:** https://github.com/denniscwylie/sarks.

**Supplementary information:** appended to document.

## 1 Introduction

Discrete sequences—of tones, of symbols, or of molecular building blocks—can provide clues to other characteristics of the entities from which they are derived: a phrase in a bird’s song can reveal which species it belongs to, the use of an idiomatic expression can pinpoint a speaker’s geographic origin, and a specific short string of nucleotide residues can illuminate the function of a DNA domain. In these examples, insights are gleaned from informative *motifs*—short subsequences that match some frequently recurring discernible pattern.

Of particular interest are DNA regions modulating differential gene expression. The regions contain motifs that produce defined patterns of gene expression, however the details of how and which motifs are needed for expression specificity remain poorly understood.

We present a broadly-applicable algorithm for identifying DNA regulatory regions that support differential gene expression. Our strategy is predicated on the following suppositions: (a) gene expression regulatory regimes involve the binding of transcription factors (TFs) to sites on non-coding DNA in the vicinity of a transcription start site (TSS) (Maston *et al.* (2006); Nguyen and D’haeseleer (2006)); (b) TFs act combinatorially to attract and repel transcription machinery (Walhout (2006)); (c) the same TF binding site may appear multiple times within a stretch of DNA, interspersed with other binding sites (Gotea *et al.* (2010)); and (d) there is more than one solution: different genes, even those co-expressed within a single cell, may rely on different regulatory mechanisms (Badis *et al.* (2009)). In accord with these suppositions, we aim to identify TF binding sites associated with enriched transcripts and scrutinize their arrangement for significant patterns that can then be evaluated experimentally.

Many motif identification methods have been described. Consensus-based methods such as Weeder (Pavesi *et al.* (2001, 2004)) focus on fixed-length motifs that repeatedly occur (with few mismatches) in sequences of interest, and can be efficiently implemented using suffix trees (Sagot (1998); Marsan and Sagot (2000); Pavesi *et al.* (2001)). Alternately, profile-based methods such as MEME (Bailey and Elkan (1995); Bailey *et al.* (2006, 2009)) build a probabilistic motif profile to be compared to a background model in order to classify subsequences as either matching the motif or not.

In contrast, discriminative methods (Sinha (2003)) identify motifs that differentiate one set of sequences (e.g., promoter regions for genes with a given expression pattern) from another (e.g., reference promoter regions). Many approaches have been applied to this differentiation problem (e.g., Segal *et al.* (2002); Segal and Sharan (2005); Redhead and Bailey (2007); Fauteux *et al.* (2008); Valen *et al.* (2009); Huggins *et al.* (2011); Yao *et al.* (2014)). One popular example, DREME (Bailey (2011)), employs Fisher’s exact test to compare counts of motif matches in the target/positive sequences with counts in the background/negative sequences. HOMER (Heinz *et al.* (2010)) uses similar hypergeometric enrichment calculations, but couples them to a zero-or-one-occurrence-per-sequence (ZOOPS) scoring approach. The recent motif finder STEME (Reid and Wernisch (2014)) extends a suffix tree-based approximate expectation-maximization approach (Reid and Wernisch (2011)) into a practical tool capable of discriminative motif discovery.

When discriminative methods are applied to differential gene expression, they impose a binary representation (such as elevated or not elevated expression). However, differential gene expression is generally described using a continuous measure (*t*-statistics, *f*-statistics, etc.), with some genes more affected than others by a difference in state. It is more useful, therefore, to use “correlative motif discovery,” which seeks motifs whose presence signals a trend towards higher or lower values of the continuous measure. A few such correlative algorithms have been described, including MOTIF REGRESSOR (Conlon *et al.* (2003)), which first applies the (non-correlative) MDScan (Liu *et al.* (2002)) algorithm to identify motifs in a subset of high-scoring sequences, then filters the motif set based on the predictive value of regression models based on the selected motifs. Another correlative algorithm, FIRE (Elemento *et al.* (2007)), iteratively optimizes the mutual information between sequence scores and occurrences of candidate motifs, starting from a set of most informative “seed” motifs.

Both of these algorithms may be seen as applying correlational information (regression or mutual information, respectively) as a filter to select and refine a set of candidate motifs generated in a non-correlative manner.

The generation of a seed motif set paves the way for sequence ranking by counting occurrences of the uncovered motifs within each sequence *ω_b_*. However, as the number of possible motifs of length *k* grows exponentially with *k*, given a fixed set of sequences *{w_b_}* and a suitably large *k*, only a fraction of possible length-k motifs will be observed in any sequence *ω_b_*. For example, in 1000 sequences *ω_1_*,…, wiooo each of length |*ω_b_|* = 1000, at most one million k-mers of any length *k* can be found.

In contrast, we aimed to develop SArKS as an algorithm for correlative motif discovery that does not require seed motifs to minimize the possibility of missing informative motifs due to suboptimal seeding. Our solution was to focus on observed substrings of the sequences *ω_b_*, not all possible k-mer patterns that could be present in the *ω_b_*.

Specifically, SArKS relies on suffixes of *ω_b_*—substrings formed by deleting the beginning of a string. As there are only |*ω_b_|* non-empty suffixes of *ω_b_*, SArKS is able to process all suffixes of its input sequences even when when they are long and/or numerous. SArKS then assesses suffix similarity by lexicographic sorting: just as words sharing a common prefix are found close together in a dictionary, suffixes starting with a shared k-mer are assigned similar numeric positions in the sorted list of all suffixes (Figure 1). By correlating sorted suffix position with suffix sequence score using kernel smoothing, SArKS develops this idea into a *de novo* motif discovery algorithm, with a natural extension for identification of longer multi-motif domains (MMDs) spanning tens to hundreds of bases (Section 2).

We applied SArKS to two RNA-seq data sets using nonparametric permutation testing to compute significance thresholds for correlative motif discovery and to estimate false positive rates. We demonstrate that SArKS outperforms existing algorithms at identifying correlative motifs in cross-validation testing scenarios. The top motif patterns and MMDs identified by SArKS include known regulatory elements (Mathelier *et al.* (2015); Elbarbary *et al.* (2016)). Thus, the correlational motif discovery approach used by SArKS takes full advantage of differential expression RNA-seq data to illuminate prospective sequence-dependent mechanisms of gene expression regulation.

## 2 Methods

Symbolic notation is described both when introduced and systematically in Section S2.1.

Given *n* sequences *ω_b_* (also referred to as words) with associated scores *yb*, the essential steps of the algorithm (illustrated in Figure 1 and described in Section 2.1) consist of:

1. concatenating all the sequences *ω_b_* into one supersequence *x;*
2. constructing the suffix array [*s_i_*] of this supersequence (Equation (2)), where *i* indexes all suffixes of *x* sorted into lexicographic order;
3. mapping suffix positions *i* back to the sequences *ω_b_i__* from which the beginnings of the associated suffixes are derived (Equation (3)); and
4. for each *i*, applying kernel smoothing to locally regress the sequence scores *yb_j_* on suffix positions *j* lexically near *i* (Equation (4)).

**Figure 1.**
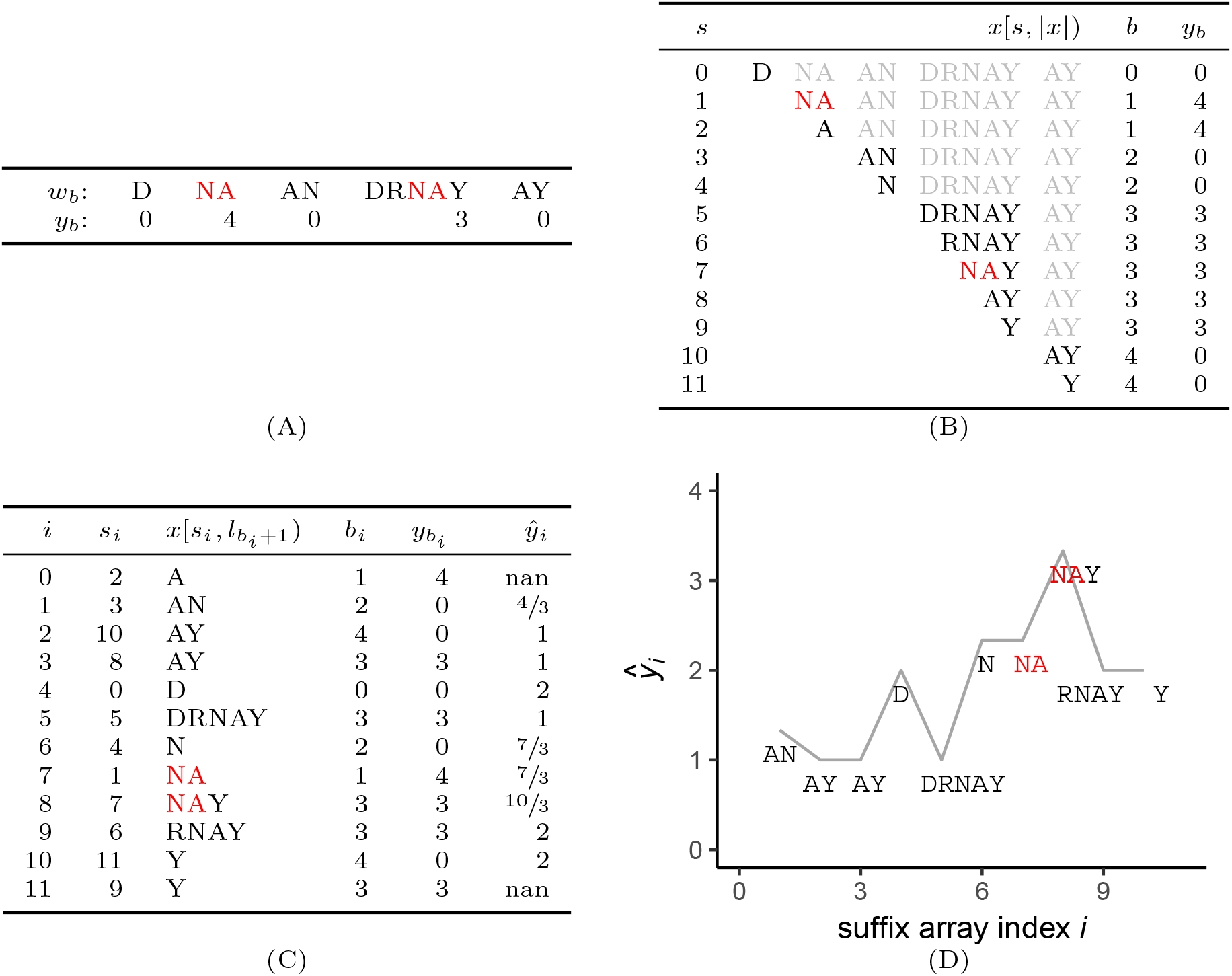
Overview of SArKS method. **(A)** Concatenation of sequences *ω_b_* (end-of-sequence character indicated by white space instead of $ for visual clarity) to form string *x* = D$NA$AN$DRNAY$AY$. (B) Table of all suffixes of *x* (part of each suffix following first end-of-sequence character shown in light gray), along with index *b* of input sequence *ω_b_* each suffix derived from and score *yb* associated with *ω_b_*. (C) Sorted suffix table indicating suffix array index *i*, suffix array value *s_i_*, suffix (sequence following first end-of-sequence character has been removed), sequence of origin *bi*, associated score *yb_i_*, and smoothed score *ŷ_i_* generated using smoothing window of size 3 (kernel half-width κ = 1). (D) Smoothed scores *ŷ_i_* plotted against suffix array index *i*, indicating peak at *i* = 8 corresponding to suffix NAY of input sequence DRNAY. Note that prefix NA of this suffix is longest substring common to the two input sequences w1 and *ω3* with scores *yb* > 0.

We thus encode the motif pattern corresponding to the first few characters of the suffix of *x* beginning at character *s_i_* with the numerical suffix array index value *i*. Because *i* gives the position of a suffix in the lexicographically sorted list of suffixes of the concatenated supersequence *x*, multiple occurrences of a highly conserved motif—even if they derive from different sequences *ω_b_*—will be consolidated into a run *i, i +* 1, …, *j* of consecutive index values. Averaging together runs of *j — i* consecutive scores by kernel smoothing using a kernel of width *j — i* thus offers a way to compare the scores *yb_i_*, *yb_i+1_, …, yb* to the overall score distribution (1).

### 2.1 Motif selection

Concatenate all words *ω_b_* (each assumed to end in the line-terminator character $ lexically prior to all other characters) to form word

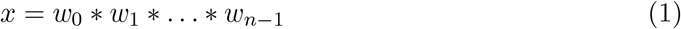

of length 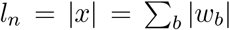. Define also 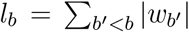. Then *x*[*l_b_*,*l_b+1_*) = *ω_b_*; that is, the substring of the concatenated string starting at position *l_b_* (inclusive) and ending immediately before position *l_b+1_* (exclusive) is the sequence *ω_b_* (we denote the first character of a string ω by ω[0], the second ω[1], etc.).

Lexically sort suffixes *x_s_* = *x*[*s*, |*x*|) into ordered set

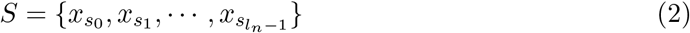

thereby defining suffix array [*s_i_*] mapping index *i* of suffix in *S* to suffix position *s* in *x* (in our software we rely on the Skew algorithm (Kärkkäinen and Sanders (2003)) modified to use a difference cover of 7 and implemented in `SeqAn` (Döring *et al.* (2008)) to efficiently compute the suffix array).

Define block array [*b_i_*] by

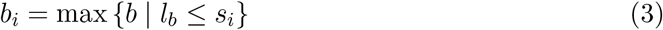

mapping index *i* of suffix in *S* to block *b* containing suffix position si. The block array then tells us that the character *x*[*s_i_*] at position *s_i_* in the concatenated string *x* is derived from *ω_bi_*[*s_i_* – *l_bi_*] in the sequence *ω_bi_*.

Calculate smoothed scores as locally weighted averages

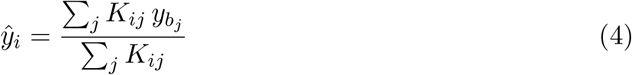

where the kernel *K_ij_* acts as a weighting factor for the contribution of the score *yb_j_* to the smoothing window centered at sorted suffix index i. *K_ij_* is used to measure how similar (the beginning of) the suffix *x*[*s_j_*, |*x*|) is to be considered to (the beginning of) the suffix *x*[*s_i_*, |*x*|) in the calculation of a representative score *ŷ_i_* averaged over suffixes similar to *x*[*s_i_*, |*x*|). As the suffixes have been sorted into lexicographic order, the magnitude of the difference *i* – *j* reflects this similarity: the key idea of the kernel smoothing approach described here is that Equation (4) with *K_ij_* defined to be a function of |i-j| may therefore offer a computationally tractable approach for identifying similar substrings which tend to occur preferentially in high scoring words *ω_b_*.

In this work we use a uniform kernel

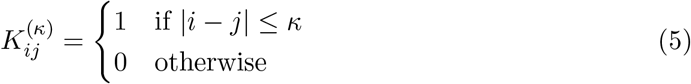

which allows Equation (4) to be computed in terms of cumulative sums:

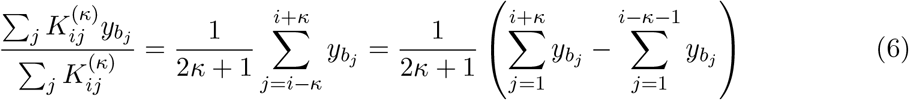

The kernel half-width κ appearing in Equation (5) is an important adjustable parameter controlling the degree of smoothing. Increasing κ smooths over more diverse suffixes, potentially increasing statistical power at the expense of the resolution of the detected motifs (i.e., length of k-mer prefix common to suffixes in the smoothing window). We recommend investigating a range of values of this parameter as is illustrated in Section 3.

Set length 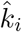 for k-mer associated with suffix array index *i* by averaging locally the length of suffix sequence identity:

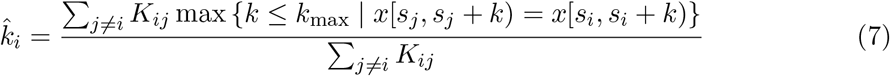

where *k_max_* functions both to increase computational efficiency and to make ki more robust in the presence of a small number of long identical substrings. All results presented here based on *k_max_* = 12: This value was selected as *k_max_* ≈ log_4_|*x*| where *x* is the longest concatenated sequence string considered in Section 3.2.1, so that for k > *k_max_* there are more distinct k-mers than there are positions for such k-mers to occur in all of the sequences *ω_b_* composing *x*.

Equation (7) is similar to Equation (4) except that: (a) Equation (7) smooths the length of the longest prefix on which the suffixes |) and |) agree instead of smoothing the score *yb*_j_ as in Equation (4); and (b) Equation (7) omits the central term *i = j* as it trivially compares the suffix beginning at *s_i_* to itself and is thus uninformative.

A straightforward approach to identifying correlative motifs using SArKS would then be to set a score threshold *θ* and take motifs to be k-mers prefixing the suffixes starting at the positions *s_i_* in the concatenated string *x*. This is the essence of our method, though below we add two filters designed to pinpoint the optimal locations *s_i_* at which to initiate motifs and, in Section S2.2, to remove likely false positive positions.

Defining the negative spatial shift operator η(*i*) which yields the unique suffix array index corresponding to the spatial position immediately prior to *s_i_*, so that *s_η_*(i) = *s_i_ —* 1, as well as the positive shift operator *ρ*(*i*) similarly defined by the condition *s_ρ_*(i) *= s_i_ +* 1, we start with a preliminary filtered suffix array index set I consisting of those *i* for which (1) the smoothed score *ŷ_i_* > *θ* and (2) *ŷ_i_* is not less than the smoothed scores of the spatial positions in *x* immediately adjacent to *s_i_* (i.e., *s_i_* must be the loci of a peak in plot of *ŷ_i_* versus spatial position *s_i_*):

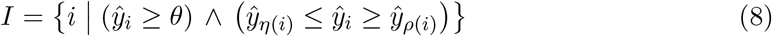

from which we obtain the associated set *M* of k-mers beginning at the positions *s_i_* in *x* by

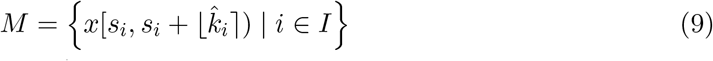

where [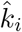] is the nearest integer to 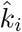. Strategies for setting the filtering threshold *θ* based on the permutation testing method are described in Section S2.5. In the next section we recommend one additional filter—designed to limit the impact of intra-sequence tandem repeats on reported motifs—to be incorporated into the definition of the index set I, replacing Equation (8) by Equation (10), for use in Equation (9).

The k-mers composing the set *M* (Equation (9)) constitute the SArKS motif set when spatial smoothing is not employed. When spatial smoothing is employed to detect multi-motif domains (Sections 2.3 and S2.4), a modified procedure for merging spatially contiguous motifs within such domains leads to Equation (S9) for the final k-mer motif set M_spatial_.

### 2.2 Limiting the impact of intra-sequence repeats

The frequent occurrence of short tandem repeats in the genome (Ellegren (2004)) can cause smoothing windows to be skewed towards a relatively small number of distinct sequences (discussed in Section S2.2). As a result, the smoothed motif scores may reflect fewer input sequences, reducing precision and increasing false positive rates among the high scoring motifs. To filter out such false positives, section S2.2 introduces the Gini impurity score *g_i_* measuring the “effective sequence count” contributing to the smoothing window centered at *i*, while Section S2.5 demonstrates that *g_i_* predicts the variance of the smoothed score for suffix array index *i* under the null hypothesis of independence between sequence and score. We can thus modify equation (8) to remove potential false positive *i* values characterized by low Gini impurities *g_i_*:

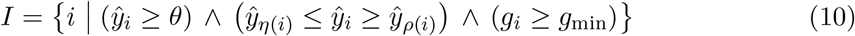

screening out positions *i* for which the repeated occurrence of a few high-scoring words in the window centered at *i* leads to *ŷ_i_* ≥ *θ*.

### 2.3 Spatial smoothing to identify multi-motif domains (MMDs)

Existing motif discovery approaches recognize the tendency of regulatory motifs to cluster into domains (Wasserman and Sandelin (2004)). Our algorithm exploits this feature, identifying candidate regulatory regions through the application of a second round of kernel-smoothing over suffix positions *s_i_* within words:

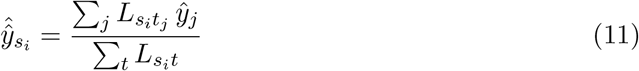

where we use uniform kernels of the form

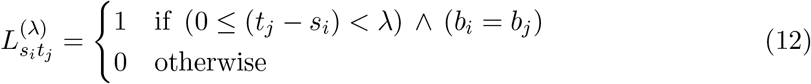

(generally with width λ ≠ *κ*) to search for regions of length λ with elevated densities of high-scoring motifs. Note that 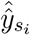 defined by Equation (11) is indexed not by suffix array index *i* but by suffix array value *s_i_* giving the spatial position *s_i_* in the concatenated word *x*. Spatial smoothing requires a threshold θ_spatial_ ≠ θ, as the doubly-smoothed scores 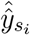 tend to be less dispersed compared to the singly-smoothed *ŷ_i_*. The threshold θ_spatial_ can be used to define an index set I_spatial_ in a manner similar to how I is defined by Equation (10). This procedure is detailed in Section S2.4, which additionally defines the set J_spatial_ of suffix array indices *i* corresponding to the starting positions of MMDs. It then details the procedure adopted by SArKS to merge spatially contiguous motifs within the same MMD, yielding the set I_spatial_ of suffix array indices *i* and merged motif lengths 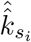 required to obtain the merged motif set M_spatial_ analogous to equation (9).

### 2.4 Permutation testing to establish significance of motif set

The significance of the correlation between the motifs uncovered by SArKS and the sequence scores *yb* can be evaluated by examining results obtained when the sequences *ω_b_* and the scores *yb* are independent of each other. To this end, the word scores *yb* are subjected to permutation π to define *yb^(π)^* = *y_π_(b)*. If the permutation π is randomly selected independently of both the sequences *ω_b_* and the scores *yb*, any true relationships between sequences and scores will be disrupted. Section S2.5 and Section S2.6 develop the strategy used by SArKS to set thresholds *θ* (and/or θ_spatial_) for each combination of parameters *κ, λ,g_min_* to control the overall false positive rate.

## 3 Results and discussion

### 3.1 Illustration of SArKS using simulated data

To illustrate SArKS, we first applied it to a simple simulated toy data set in which 30 random sequences *ω_b_* were generated with each letter *ω_b_*[*s*] drawn independently from a Unif {A,C,G,T} distribution. We then embedded the k-mer motif CATACTGAGA (*k* = 10) in the last 10 sequences (i.e., those *ω_b_* with *b ≥* 20) by choosing a position *sb* independently for each sequence *ω_b_* from Unif {0,…, |*ω_b_| — k}* and replacing *ω_b_*[*sb, sb* + *k*) with the motif. Scores were assigned to the sequences according to whether the motif had been embedded: *yb* = 0 if *b* ∊ [0,20), *yb* = 1 if *b ≥* 20.

The kernel half-width *κ* = 4 was used to obtain smoothing windows of size similar to the number of motif-positive sequences, *2κ +* 1 ≈ |{*b* | *yb* = 1}|. As this number cannot be known in advance when applying SArKS to real data, in practice we recommend testing a range of *κ* values as done in Section 3.2.1 below.

Figure 2 plots *ŷ_i_* as obtained from Equation (4) applying the method of Section 2.1 to search for motifs. The highest peaks in the plot correspond to the positions of various substrings of the embedded motif, and correspond to the set *M* of k-mers defined by the *x*[*s_i_, s_i_* + |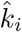|) column of Table 1.

Removing nested motifs from Table 1 as described in Section S2.3, Equation (S7) leaves only the rows for *i* ∊ {2257,2258,2256,1462,1458,1463}. Applying Equation (S8) then extends the 8-mer ATACTGAG of the rows *i* ∊ {1462,1458,1463} to the full 10-mer, so that, following Equation (S9), the final k-mer set *M’* = {CATACTGAGA} is recovered (Table 1).

Section S2.5 illustrates the utility of setting a minimum Gini impurity *g_min_* during motif selection to reduce the false positive rate: 190 out of 1000 random permutations generated at least one position *i*^(π^) for which 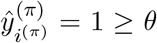 (for this toy model *θ* was taken to have the maximum possible value of 1), but only 20 of these permutations yield any results if g_min_ = 0.8506 (following Equation (S5) with γ = 0.1) is applied. Based on these results, we can derive a 95% confidence interval of (1.2%, 3.1%) for the family-wise error rate (FWER, a type of false positive rate; see Section S2.6).

### 3.2 Uncovering promoter motifs associated with differential gene expression

We set out to analyze two published RNA-seq data sets (Mo *et al.* (2015); Close *et al.* (2017)) using SArKS. The first study presented transcriptome data for adult mouse neurons sorted according to cell class (Mo *et al.* (2015)). In particular, this study was among the first to profile parvalbumin-expressing (PV) interneurons, a major inhibitory subclass in the mammalian neocortex. PV basket and chandelier neurons are intimately involved in the microcircuitry of sensory processing, memory formation and critical period plasticity (Cobb *et al.* (1995); Klausberger and Somogyi (2008)). Dysfunction of PV interneurons has been linked to autism and schizophrenia (Lewis *et al.* (2005)), and the ability to access these neurons using a cell-type specific promoter has been a priority for brain scientists. The second study examined transcriptomes of differentiating interneurons at several developmental time points (Close *et al.* (2017)). In the sections below, we describe the parameters and results of SArKS analyses for both data sets. In Section S3.2.2, we inspect the SArKS-elicited motifs associated with PV neurons and demonstrate how to extend the application of SArKS to identify promoter multi-motif domains (MMDs).

**Figure 2.**
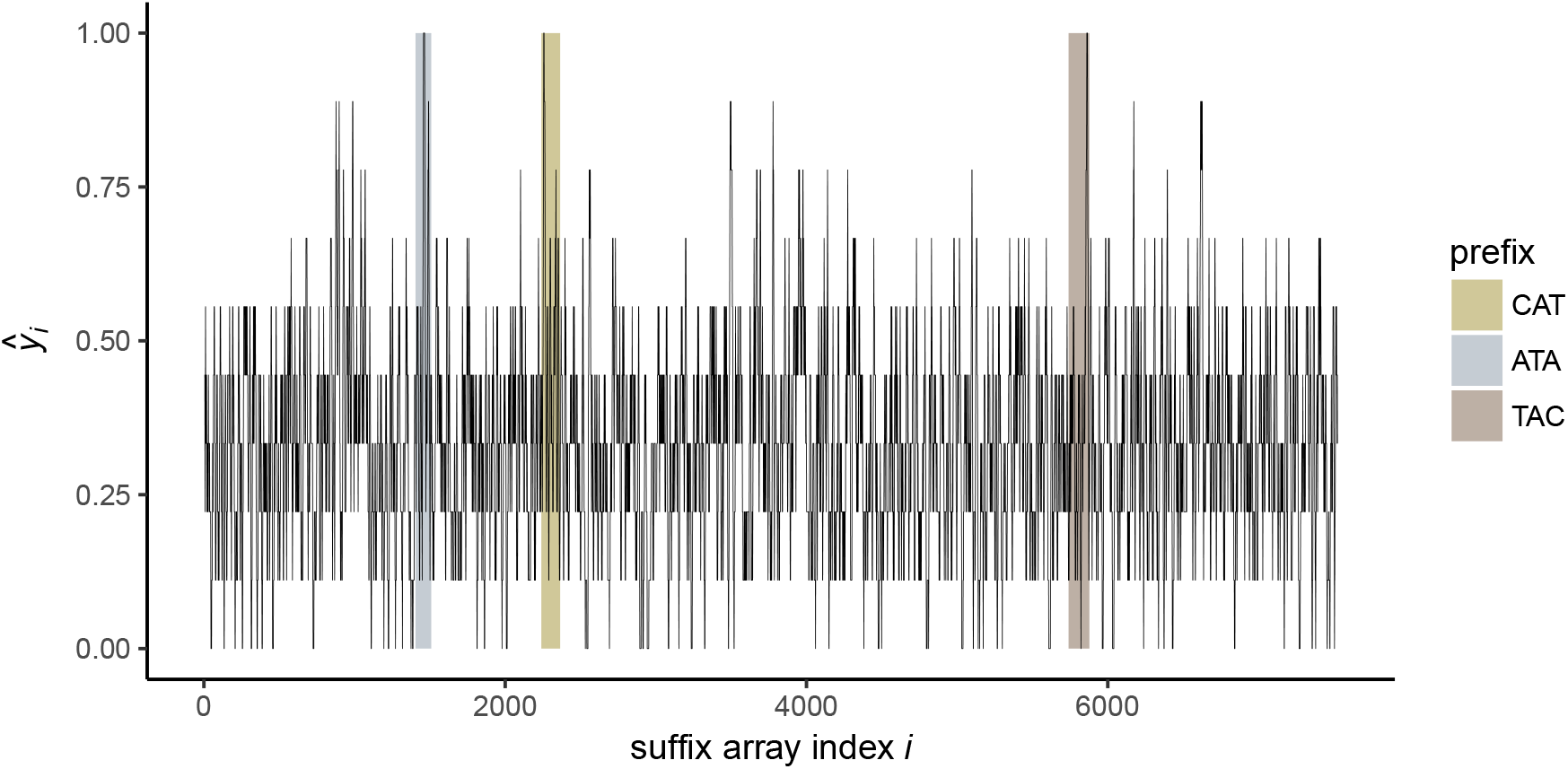
Locating peaks in kernel-smoothed scores iˆ. Kernel-smoothed scores *ŷ_i_* (Equation (4), using kernel half-width *κ* = 4) are plotted against suffix array index *i* for simulated data set. Gold, silver, and bronze bars indicate positions in lexicographically sorted table of suffixes beginning with prefixes CAT, ATA, and TAC, which correspond to the first five characters of embedded motif CATACTGAGA. Detailed information on the peak locations at which the smoothed score *ŷ_i_* = *1* is presented in Table 1 below.

#### 3.2.1 Data set 1: cell class-specific transcriptome analysis

We analyzed RNA-seq gene expression data from mouse neocortical neurons pooled based on genetically defined cell classes (Mo *et al.* (2015)) to identify regulatory motifs associated with parvalbumin (PV) neuron-specific gene expression.

After accounting for differential expression and chromatin accessibility (Section S2.7.1), we examined two sets of sequences for 6,326 unique transcripts. The first set covered upstream regions −3000 base pairs (bp) to the transcription start site (TSS), the second set extended from the TSS to +1000 bp.

We tested a range of half-window sizes *κ* ∈ {250, 500,1000, 2500} with the maximum value of 2500 selected to produce a smoothed window of size *2κ +* 1 = 5001 similar to the number of input sequences (6326). Note that smaller windows are less likely to contain sample multiple suffixes from the same promoter sequence: in particular, windows of width greater than the number of distinct sequences must contain multiple suffixes from at least one promoter sequence.

For each half-window size *κ*, we applied two minimum Gini impurity values *g_min_* set according to Equation (S5) with first 7 = 0.1 and then 7 = 0.2. Also for each value of *n*, we examined three separate spatial window sizes λ ∈ {0,10,100}. These values were selected to investigate the performance of SArKS using no spatial smoothing (λ = 0), using a window λ = 10 of the typical length scale of eukaryotic transcription factor binding sites (Stewart *et al.* (2012)), and using a window λ = 100 to target the low end of the enhancer length distribution (Loots (2008)). Thresholds *θ* (for analyses with no spatial smoothing) or *θ*_spatial_ (for analyses with λ ∈ {10,100}) were set according to the permutation testing strategy detailed in Section S2.5 using *R* = 100 permutations.

**Table 1.**
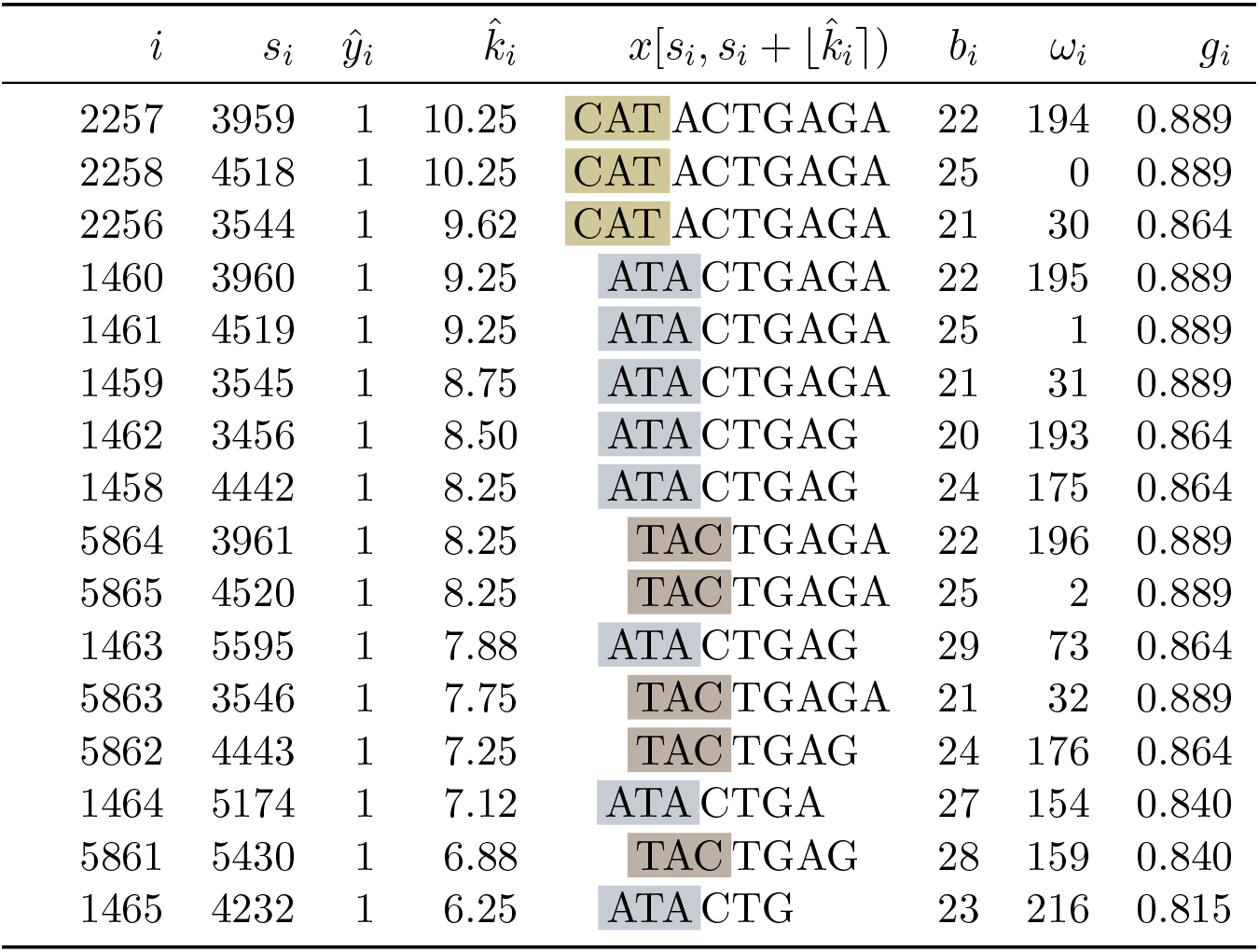
Suffix array peak positions with *ŷ_i_* > *θ*. Illustration of motif selection process (Section 2.1) applied to simulated data (using kernel half-width κ = 4). All positions for which sequence smoothed score *ŷ_i_* > *θ* = 1 are shown; table is sorted in descending order of the estimated motif length k_i_. Columns indicate values of key variables for the suffix associated with the corresponding peak: (*i*) suffix array index *i* giving position of suffix in lexicographically sorted list of all suffixes; (*s_i_*) suffix array value *s_i_* giving spatial position of suffix in concatenated sequence *x;* (*ŷ_i_*) kernel smoothed score *ŷ_i_* (Equation (4)); (k_i_) estimated length 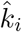 (Equation (7)) of conserved |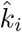|-mer prefix of suffixes within smoothing window centered on suffix array index *i;* (*x*[*s_i_, s_i_ +* |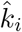])) the corresponding conserved |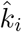|-mer *x*[*s_i_*, *s_i_* + |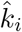|) (Equation (9)); (*bi*) the input sequence *bi* (Equation (3)) from which the suffix is derived; (ω*i*) the spatial position ω*i* at which the suffix is found within sequence *bi;* and (*g_i_*) the Gini impurity *g_i_* (Equation (S4)) for the smoothing window centered at *i*. Note that each of these peaks corresponds to a suffix derived from a position within the first three characters of an instance of the embedded motif CATACTGAGA. Gold highlighting indicates peaks starting from the first character of the embedded motif, silver the second, and bronze the third.

| $i$ | $s_i$ | $\hat{y}_i$ | $\hat{k}_i$ | $x[s_i, s_i + \lfloor \hat{k}_i \rfloor]$ | $b_i$ | $\omega_i$ | $g_i$ |
| --- | --- | --- | --- | --- | --- | --- | --- |
| 2257 | 3959 | 1 | 10.25 | CAT ACTGAGA | 22 | 194 | 0.889 |
| 2258 | 4518 | 1 | 10.25 | CAT ACTGAGA | 25 | 0 | 0.889 |
| 2256 | 3544 | 1 | 9.62 | CAT ACTGAGA | 21 | 30 | 0.864 |
| 1460 | 3960 | 1 | 9.25 | ATA CTGAGA | 22 | 195 | 0.889 |
| 1461 | 4519 | 1 | 9.25 | ATA CTGAGA | 25 | 1 | 0.889 |
| 1459 | 3545 | 1 | 8.75 | ATA CTGAGA | 21 | 31 | 0.889 |
| 1462 | 3456 | 1 | 8.50 | ATA CTGAG | 20 | 193 | 0.864 |
| 1458 | 4442 | 1 | 8.25 | ATA CTGAG | 24 | 175 | 0.864 |
| 5864 | 3961 | 1 | 8.25 | TAC TGAGA | 22 | 196 | 0.889 |
| 5865 | 4520 | 1 | 8.25 | TAC TGAGA | 25 | 2 | 0.889 |
| 1463 | 5595 | 1 | 7.88 | ATA CTGAG | 29 | 73 | 0.864 |
| 5863 | 3546 | 1 | 7.75 | TAC TGAGA | 21 | 32 | 0.889 |
| 5862 | 4443 | 1 | 7.25 | TAC TGAG | 24 | 176 | 0.864 |
| 1464 | 5174 | 1 | 7.12 | ATA CTGA | 27 | 154 | 0.840 |
| 5861 | 5430 | 1 | 6.88 | TAC TGAG | 28 | 159 | 0.840 |
| 1465 | 4232 | 1 | 6.25 | ATA CTG | 23 | 216 | 0.815 |

#### 3.2.2 Data set 2: differentiating interneuron transcriptome analysis

We examined RNA-seq data for differentiating human interneurons (Close *et al.* (2017)), applying SArKS to identify promoter motifs associated with elevated gene expression in doublecortin-positive (DCX+) GABAergic neurons compared to DCX- cells. Differential expression was assessed for 6,939 genes as detailed in Section S2.7.2 and we analyzed upstream sequences (from −3000 bp to the TSS) and downstream sequences (from the TSS to +1,000 bp) as described in Section 3.2.1.

SArKS analysis was conducted using all combinations of half-window size *κ* ∈ {250,500,1000,2500} and spatial smoothing window λ ∈ {0,10,100} for the reasons described in 3.2.1. However, for this data set, the minimum Gini impurity thresholds were computed using only γ = 0.1— we had seen little benefit from including the higher value γ = 0.2 in our experience with the Mo 2015 data set (see Section 3.2.4). Thresholds *θ* or θ_spatial_ were set according to the permutation testing strategy detailed in Section S2.5 using *R* = 100 permutations.

#### 3.2.3 Benchmark comparisons for correlative motif discovery

We conducted a cross-validation benchmarking study to compare SArKS correlative motif discovery performance to that of five motif search algorithms. Two of these methods, FIRE (Elemento *et al.* (2007)) and MOTIF REGRESSOR (Conlon *et al.* (2003)), as they rely on alternative approaches to correlative motif discovery. The remaining algorithms, DREME (Bailey (2011)), HOMER (Heinz *et al.* (2010)), and STEME (Reid and Wernisch (2014)) are popular discriminative methods which we have run by discretizing our score data with promoter sequences *b* considered ‘positive’ sequences if the score *yb* ≥ 2, ‘negative’ otherwise. While there is a definite loss of information in this discretization—the avoidance of which is one of the primary motivations for the introduction of SArKS, as well as other correlative motif algorithms—we were interested in direct comparison of correlative and discriminative algorithms to assess the degree to which correlative algorithms actually benefit from avoiding discretization.

We split the 6,326 transcripts selected from the Mo 2015 data set into five disjoint subsets *V_1_, V_2_, …*, V_5_ (the name *V_f_* intended to suggest the f^th^ validation set) of approximately equal size (| V_1_| = 1,266, while |*V_f_*| = 1, 265 for f > 1). The set of 6,939 genes selected from the Close 2017 data set was similarly partitioned into disjoint cross-validation folds.

For both data sets, for both promoter sequence ranges investigated, and for each of the algorithms evaluated, five separate motif identification analyses were conducted corresponding to the five cross-validation folds *V_f_*. For each analysis f ∈ {1,…, 5}, motif discovery was performed using the sequences and scores from all folds except *V_f_*: the genes assigned to *V_f_* were instead held out for validation of the discovered motif (so that the set of genes used to learn the motif sets was in each case disjoint from the set of genes used for validation). Existing algorithms were run at their default parameter settings where possible; exact specifications are given in Section S2.8, while SArKS parameters were set as described in Sections 3.2.1-3.2.2.

We used `tomtom` (Gupta *et al.* (2007)) to compare the pooled motif sets identified by each algorithm. Figure S3 shows the resulting overlap between motifs sets by algorithm: for each of the benchmarked algorithms, the majority of identified motifs had a SArKS-identified counterpart. SArKS also identified many additional motifs.

The Pearson correlation between the count of occurrences of a given motif in sequence *ω_b_* with the score *yb* across the sequence-score pairs (*ω_b_, yb*) provides a natural metric for assessing correlative motif discovery performance. Figure 3 plots the estimated Pearson correlation values for each motif identified (by each algorithm) evaluated using the held-out validation set *{*(*ω_b_, yb*) | *b* ∈ *V_f_*} appropriate for the fold f in which the motif was discovered (with 3A and 3B presenting results for the Mo 2015 and Close 2017 data sets, respectively).

As Figure 3A-B and Figure S3 demonstrate, the number of motifs identified by different algorithms can be highly variable: DREME, FIRE <C HOMER, MOTIF REGRESSOR ¡ SArKS (the motif count for STEME is a fixed input parameter). The interpretation of the number of motifs is, however, complicated by two factors: (1) the occurrence rate of individual motifs in the relevant biological sequences (promoters, etc.) may differ substantially (e.g., longer motifs may occur less frequently) and (2) some motifs may be very similar in sequence.

**Figure 3.**
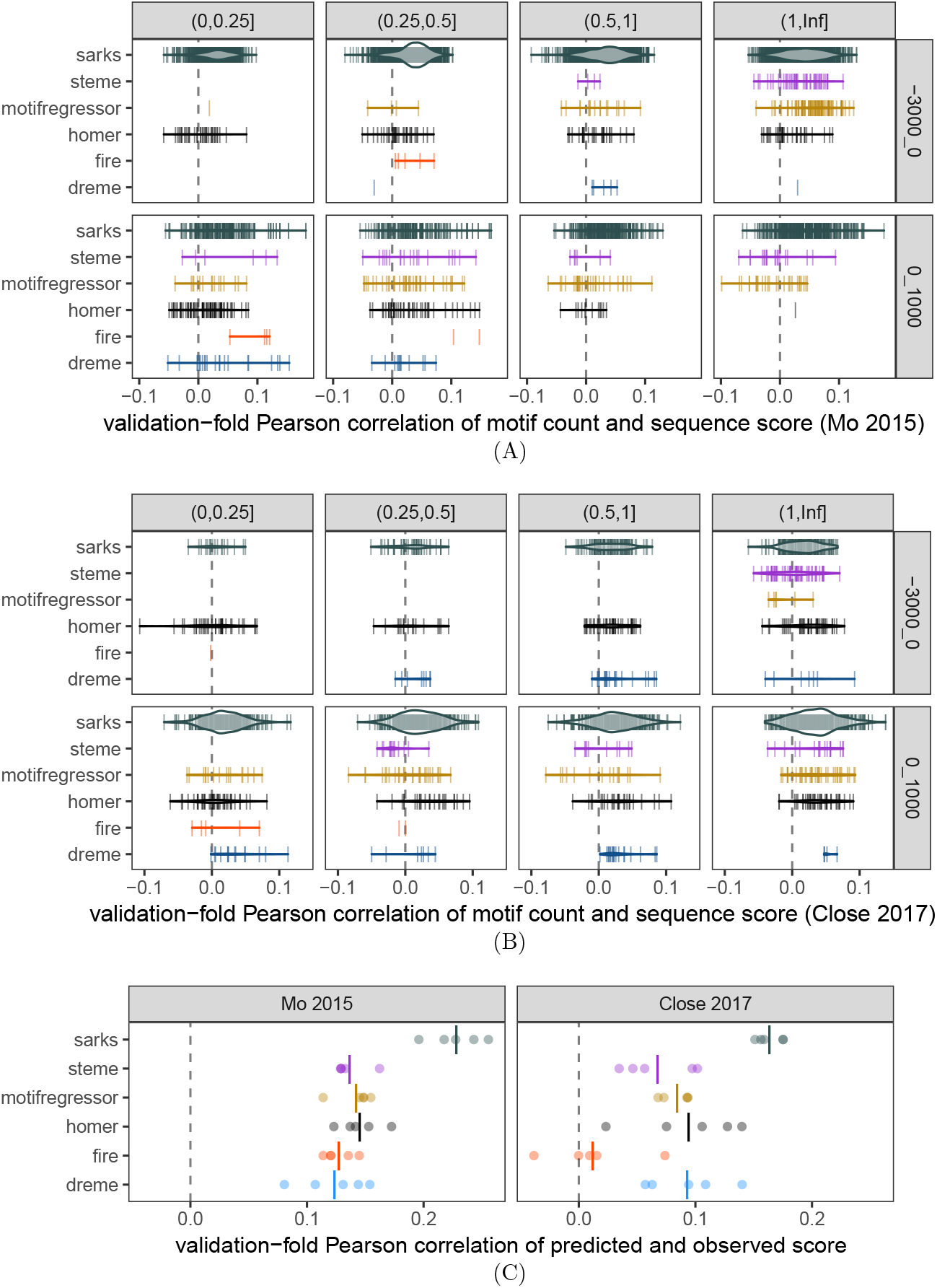
Benchmark comparisons of correlations between motif counts and gene specificity scores in held-out validation subsamples. **(A)** Each vertical line represents a motif identified by the indicated algorithm in one of the five cross-validation folds for the Mo 2015 data set (Mo *et al.* (2015)). The horizontal position of the line encodes the Pearson correlation coefficient of the motif count with the associated sequence score (calculated using only the genes in the held-out validation set for the cross-validation fold in which the motif was identified). The count for a given motif in sequence *ω_b_* was assessed using fimo (Grant *et al.* (2011)) for DREME, HOMER, MOTIF REGRESSOR, and STEME—all of which represent motifs as position-weight matrices—and using a simple regular expression search for FIRE (which returns regular expression representations of motifs) and for SArKS k-mers. In all cases, motif counts were based on motif occurrences on either the forward or reverse strand. Row: sequence region for motif counts, either 3kb upstream or 1kb downstream of TSS; column: interval containing average number of occurrences of motif within sequence region across all analyzed genes. Widths of violins represent motif density and are scaled consistently across all panels. **(B)** Same as (A), except applied to Close 2017 data set (Close *et al.* (2017)). **(C)** Motif regression model predictions correlate with gene specificity scores in held-out cross validation subsamples. Each of five cross-validation folds is plotted as separate point for each algorithm. Each regression model was built using feature vector constructed by concatenating counts of upstream motifs in upstream regions with counts of downstream motifs in downstream regions. Left panel: results of modeling applied to Mo 2015 data set; right panel: same for Close 2017 data set. Vertical lines indicate mean Pearson correlation across all folds.

The first of these complications is illustrated in Figure 3A-B by faceting horizontally on motif occurrence rate (count per sequence): one visible trend here is that the DREME-, FIRE-, and HOMER-identified motifs tend to occur less frequently than do the MOTIF REGRESSOR and STEME motifs, indicating that DREME, FIRE, and HOMER tend to define motifs more granularly than do MOTIF REGRESSOR or STEME. SArKS-identified motifs are spread across a wide range of per-sequence occurrence rates in this plot, as SArKS identifies both more and less granular motifs as the size of the smoothing window κ is varied through the ranges specified in Sections 3.2.1-3.2.2.

The second complication—the similarities among identified motifs—may be addressed by noting that correlative motif discovery can also be viewed as a form of feature extraction. In this vein, we can assess the performance of such algorithms by using the selected motifs as predictors to build regression models for associated sequence score *yb* based on the motif counts in the sequence *ω_b_*. Figure 3C plots validation set-estimated Pearson correlations of the predictions made by building a linear ridge regression model (using generalized cross-validation (Golub *et al.* (1979)) to select the L2 regularization parameter) with the sequence scores for each cross-validation fold by algorithm. Motifs were counted only within the sequence range in which they were identified, with these counts then merged into a single feature vector per gene to allow the regression models to consider both up- and downstream motifs simultaneously. This approach collapses the variation in quantity and quality of individual motifs down to variation of a single quantity—the regression model predictions—thereby facilitating a head-to-head comparison of motif discovery algorithms bypassing both of the complications discussed above. As the similarity of some identified motifs manifests as collinearity of regression predictors, regularization is a key component of this modeling approach.

SArKS yields better results than the other algorithms for both validation data sets (Figure 3C); aside from SArKS, the other two correlational motif discovery algorithms (FIRE and MOTIF REGRESSOR) do not appear to show a consistent advantage in performance relative to the discriminative algorithms.

If, instead of using the merged motif feature set, the regression models are built using only upstream or downstream motif counts, the results shown in Figure S2 are obtained, making clear that all six algorithms generally perform better when searching the downstream regions (for which SArKS shows a particularly strong advantage in both data sets).

Considering the downstream motif results, we noted that for every algorithm applied to the Mo 2015 data set, the motif with the highest Pearson correlation coefficient between occurrence count and parvalbumin specificity score in the held-out cross-validation fold exhibited significant `tomtom` similarity (*q* ≤ 0.1) to the ESRRA/ESRRB/ESRRG trio of TF-binding motifs documented in the JASPAR database (Mathelier *et al.* (2015)). Looking at the Close 2017 downstream motif results, we observed that the most highly cross-validation-correlated motifs for three of the algorithms—FIRE, HOMER, and SArKS—were significantly similar to all of the JASPAR motifs TGIF1/TGIF2/MEIS2/MEIS3/PKNOX1/PKNOX2.

In contrast, the top upstream motif results showed no such convergence on common JASPAR profiles: applied to the Mo 2015 data set, only one pairwise combination of two algorithms—FIRE and STEME—produced top upstream motifs (ranked by cross-validated Pearson correlation) that showed significant `tomtom` similarity (*q* ≤ 0.1) to a common JAS-PAR profile (NR5A2, whose binding motif closely resembles the ESRRA/ESRRB/ESRRG pattern mentioned above). Applied to the Close 2017 data set, no two algorithms produce motifs similar to the same JASPAR profile.

**Figure 4.**
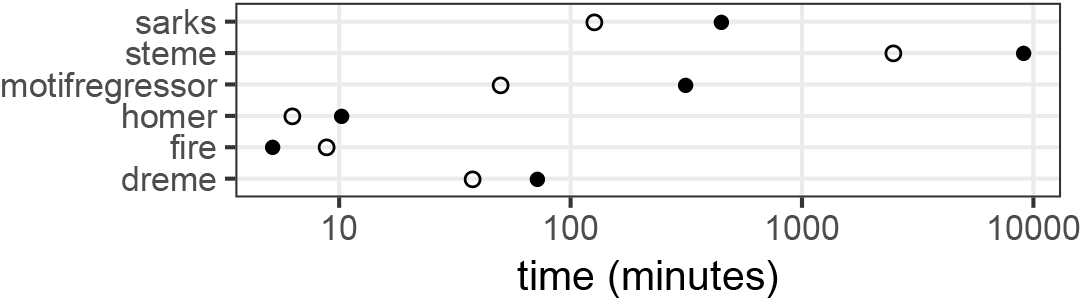
Benchmarked algorithm run times. Average run times per cross-validation fold for each motif discovery algorithm applied to either upstream (solid circles) or downstream (open circles) regions for selected genes from Close 2017 data set (for which all analyses were run on the same computer system).

We see that in those cases where all of the algorithms performed better in the cross-validation testing (downstream), the top motifs were more likely to converge on known TF-binding motifs. Interestingly, SArKS outperformed the other algorithms to a greater degree in the analyses of downstream regions than of upstream regions.

Comparison of all of the motifs discovered by the various algorithms with known TF-binding motifs is further explored in Section S3.2.1.

Finally, we compared the average run times for each of the benchmarked algorithms applied to the upstream and downstream cross-validation analyses. As is shown in Figure 4, SArKS took longer than most of the other algorithms with the exception of STEME; FIRE and HOMER are quite fast relative to the others. Further discussion of the computational complexity of SArKS is provided in Section S3.3.

#### 3.2.4 Permutational analysis of SArKS results

The permutation testing procedure used to set SArKS score thresholds can be used for directly assessing the statistical significance of the motif set SArKS reports as well. This is done by (1) following the procedure laid out in Sections S2.5 and S2.6 using a set of R randomly drawn permutations of the input sequence scores to determine threshold values for motif selection and (2) independently drawing a second set of R2 permutations from which the false positive rate corresponding to these thresholds can be estimated according to (S26).

To demonstrate this procedure, we re-applied SArKS to both the Mo 2015 and Close 2017 data sets here including all 6,326 or 6,939 selected genes (respectively) without cross-validation subsetting. We again investigated all combinations (κ, λ) ∊ {250,500,1000,2500} *x* {0,10,100} for the smoothing half-width κ and spatial length λ, computing *g*_min_ for each value of κ following Equation (S5) using the γ values indicated in Sections 3.2.1-3.2.2, and determining significance thresholds using R = 100 randomly generated permutations.

For the Mo 2015 data set, the analyses performed using the stricter *g*_min_ values obtained using γ = 0.1 yielded larger k-mer motif sets: 3,393 total distinct k-mers versus only 1,232 using γ = 0.2 for the upstream sequence set; 380 distinct k-mers using γ = 0.1 versus just 180 using γ = 0.2 for the downstream sequence set. More than 98% of the k-mers discovered using 7 = 0.2 were also identified using 7 = 0.1 (for both sequence ranges: 1,208 of the 1,232 upstream; 179 of the 180 downstream). Based on these results for the Mo 2015 analysis, we focused exclusively on 7 = 0.1 for the Close 2017 analysis, as described in Section 3.2.2.

The results above demonstrate that restrictive values of 7 can yield larger motif sets that include almost all of the motifs obtained using more permissive 7 values. This highlights the importance of the Gini impurity filter in focusing SArKS on potential motifs that appear within sufficiently many distinct sequences WB to achieve reasonable statistical confidence.

We assessed the statistical significance of these SArKS results following the method of Section S2.6 with thresholds 9 and Spatial set by Equation (S24) and Equation (S25) using z = 4. Upstream sequence analysis of the Mo 2015 set considering R2 = 250 independent random permutations resulted in 12 (4.8%) for which any of the parameter sets (ft, *g*_min_, A) yielded a nonempty set of identified motifs; for the Close 2017 set, the same procedure resulted in 8 (3.2%) nonempty motif sets. These upstream sequence results correspond to a 95% family-wise error rate confidence interval (FWER CI) of (2.5%, 8.2%) in the Mo 2015 analysis and (1.4%, 6.2%) in the Close 2017 analysis.

For the downstream sequence analysis, R2 = 250 independent permutations yielded 8 (3.2%) instances of nonempty motif sets for Mo 2015 and 1 (0.4%) nonempty motif set for Close 2017, from which we estimate 95% FWER CIs of (1.4%, 6.2%) for Mo 2015 and (0.01%, 2.2%) for Close 2017.

The role of the parameter z in Eqs (S24)-(S25) in balancing FWER against sensitivity can be seen in the analyses presented here by considering the consequence of increasing z: at z = 5 for the same 250 permutations, the permutation analysis using upstream regions resulted in nonempty motif sets in only 2 permutations for Mo 2015 or 1 permutation for Close 2017. Similarly, for the downstream regions, permutation analysis with z = 5 resulted in 4 or 1 permutation(s) respectively. The cost of these decreased false positive rates to sensitivity is apparent in that at most half of the motif fc-mers identified using z = 4 were still discovered using z = 5 in each of the analyses; for the Close 2017 upstream analysis conducted with z = 5, SArKS returned no significant motif results at all. Here we were willing to accept the FWER values associated with z = 4 (point estimates ranging from 0.4% to 4.8% in these analyses) in order to maintain a higher sensitivity.

Selection of the parameter z to appropriately balance sensitivity against false positive rate will generally depend on the range of k, A, and *g*_min_ values investigated. When SArKS analyses are conducted for many combinations of these parameters there will be correspondingly more possible opportunities for false positives, requiring a higher value of z to maintain confidence in the results. In cases where the size of the returned motif set may be large, there is an additional factor to consider: the smaller motif sets associated larger values of z benefit not only from greater statistical confidence but also from a reduction in the computational effort required to refine and process the motif set (Section S3.3).

## 4 Conclusions

We introduce SArKS as a method for *de novo* correlative motif discovery. SArKS avoids the dichotomization—and consequent loss of information (Fedorov *et al.* (2009))—of sequence scores into discrete groups as required by discriminative motif discovery algorithms. SArKS does not require specification of parametric background sequence models, instead using nonparametric permutation methods (Ernst (2004)) to set thresholds for motif identification and to estimate false positive rates. SArKS smooths over spatial motif location to identify multi-motif domains (MMDs), which can in turn help refine the identified motifs. We have benchmarked SArKS against several existing discriminative and correlative algorithms using previously published RNA-seq data: SArKS uncovered particularly large motif sets and SArKS motif sets functioned as more predictive feature sets in a cross-validated regression modeling approach than did motif sets generated by existing algorithms. SArKS thus offers an approach to motif discovery capable of fulling exploit differential gene expression data.

## Acknowledgements

The authors thank Dr. Becca Young, Eric Brenner, Brian Gereke and Dr. Preeti Mehta for helpful discussions and Dr. Ila Fiete for a critical reading of the manuscript. This project has been supported by NIH BRAIN Initiative award U01NS094330 to BVZ.

## S2 Methods

### S2.1 Notation glossary

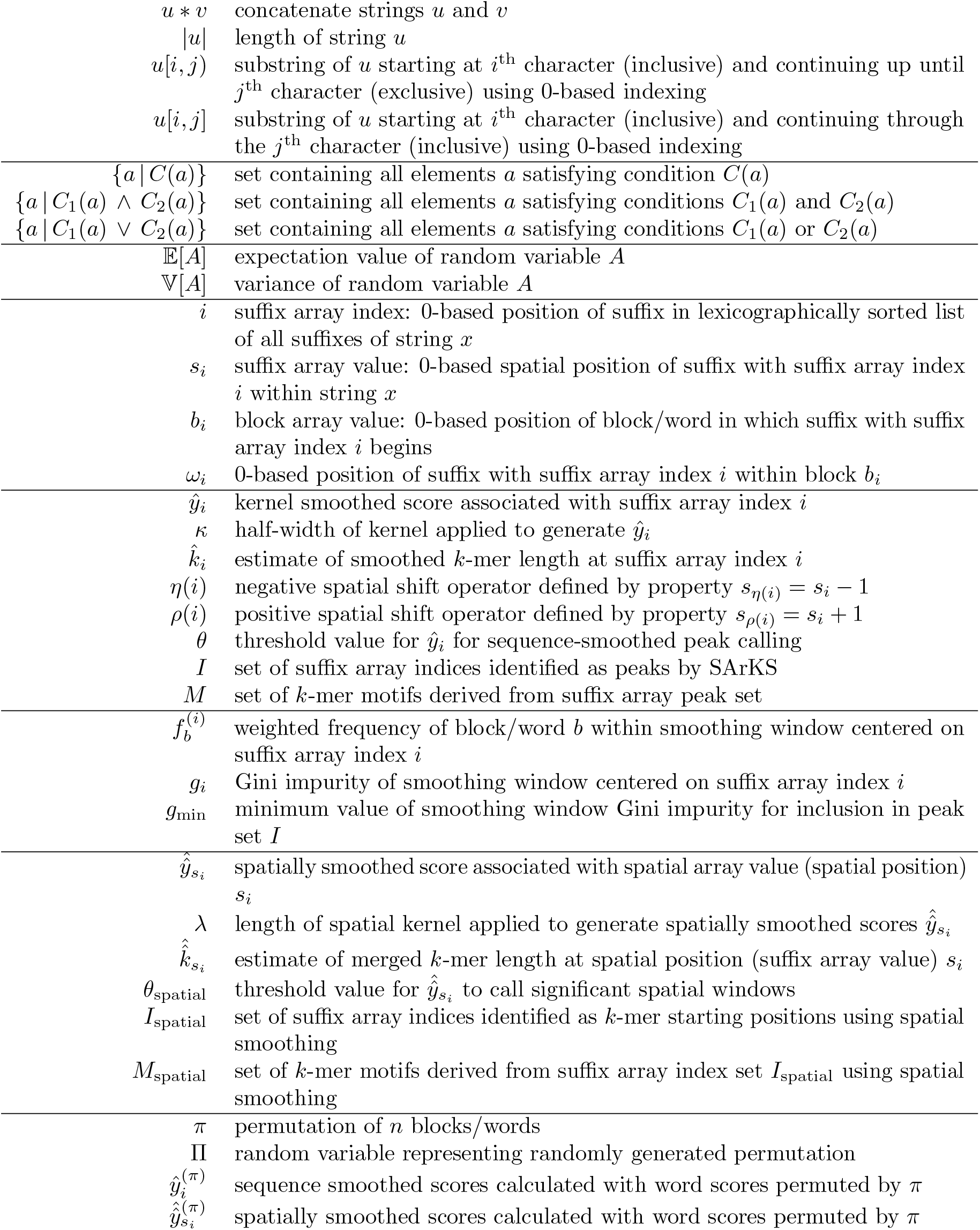

### S2.2 Limiting the impact of intra-sequence repeats

One complicating factor in the strategy described in Section 2.1 is the presence of tandem repeats (common in eukaryotic DNA (Ellegren, 2004)): if the substring *x*[*s_i_, s_i_ + rm*) (assumed to derive wholly from the single word *ω_b_*) consists of r 3> 1 repeats of the same m-mer,

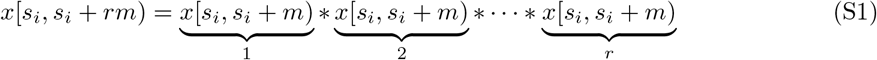

then it is likely that the sorted suffix array index positions *j* and *k* implicitly defined by *s_j_* = *s_i_ + am* and s_k_ = s_i_ + *bm* for small a, *b ≥* 0 will be close by, since, assuming without loss of generality that *a < b*,

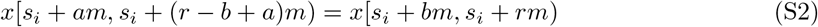

showing that the suffixes of *x* beginning at positions (si + am) and (si + bm) agree on their first (r — b)m characters. Since all of the positions *s_i_* + am for small a must come from the same word block bi they must have the same associated score *yb*_i_. If this score *yb*_i_ is particularly high, this phenomenon may lead to windows of high *ŷ_j_* values centered on *j* satisfying *s_j_* = *s_i_* + am which result from a very small number of different repeat-containing words (perhaps as few as one if the number of repeats is high enough within a single high-scoring word). We thus here develop a natural method for filtering the peak index set I to selectively remove suffix array index values *i* where the smoothing window is dominated by a few heavily repeated words *ω_b_*. The distribution of weighted word frequencies

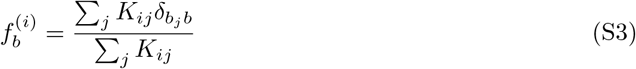

contributing to the window centered at position *i* of the suffix array table across the full word set ω may for these purposes be summarized by the associated Gini impurity (often used in fitting classification and regression trees (Breiman *et al.*, 1984)):

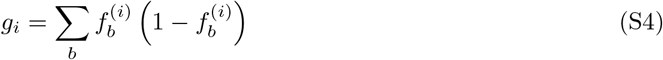

which provides a measure ranging from 0 to 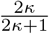 of the degree of uniqueness of the words contributing to the calculation of yi.

F (0

As a concrete example, *i* all of the weighted frequencies word frequencies fb = 1 are the same for a set of exactly *q* words *ω_b_* appearing in the smoothing window centered on i, = 1 — 1. This *g_i_ q* suggests an intuitive interpretation of (1 — gi) as the multiplicative inverse of the “effective word count” contributing to the smoothing window around i.

Section S2.5 further demonstrates that (1 — gi) is also approximately proportional to the variation of the smoothed scores yi that would be expected if there were no association between the sequences *ω_b_* and the scores *yb* (see Equation (S21) below). This proportionality suggests a simple method for selection of a *g*_min_ value at which most suffix array indices *i* will be retained while filtering out only those most likely to yield false positive results under permutation testing:

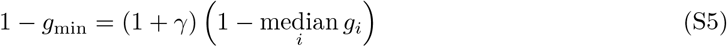

As shown in Equation (S21), setting *g*_min_ to satisfy Equation (S5) removes suffix indices *i* for which the variance of the permuted smoothed scores is greater than (1+γ) times the median value. Thus any value γ > 0 will retain the majority of positions *i* for further consideration. We have used γ = 0.1 or γ = 0.2 for all of the examples in the present work, retaining positions for which the permuted score variance is less than 110% or 120%, respectively, of the median.

**Figure S1.**
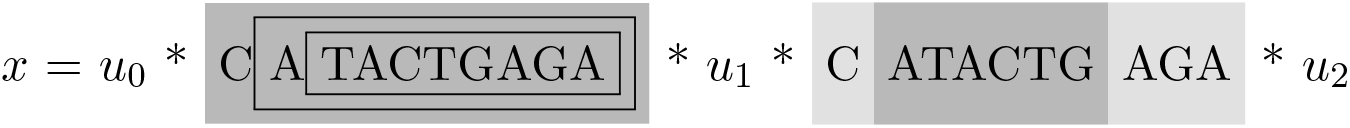
Example k-mers to be removed or extended to reduce redundancy in reported motif set. Two identified k-mers (u=CATACTGAGA on the left and v=ATACTG on the right) are indicated by the dark gray highlighting, with two additional separately identified k-mers that are part of *u* indicated within the nested black boxes. The two nested k-mers contained within the boxes inside of *u* will be removed from the discovered k-mer set by the method of Section S2.3.1, while k-mer *v* = ATACTG will be extended by the method of Section S2.3.2 to include the characters highlighted in light gray, replacing *v* with CATACTGAGA. The four k-mers indicated in this figure correspond to positions *s_i_* ∈ {3959,3960,3961,4232} from Section 3.1.

Requiring *g_i_*_*g*_min_ results in rede_ning the peak index set I to

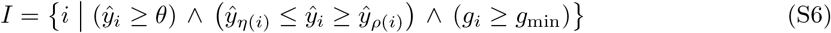

screening out positions *i* for which the repeated occurrence of a few high-scoring words in the window centered at *i* leads to *ŷ_i_* > *θ*.

### S2.3 Reducing redundancy in reported motif set

The presence of a k-mer *x*[*s_i_, s_i_ + k*) associated with a high smoothed score *ŷ_i_* can also result in high smoothed scores *ŷ_j_* when *s_j_* = *s_i_* + *m* if the substring (k — m)-mers *x*[*s_i_ + m, s_i_ + k*) also also preferentially found in higher-scoring sequences, as pictured in Figure S1. The following two steps may be added to the algorithm described in Section 2.1 in order to reduce the reporting of such substrings when they are present only as part of the full k-mer.

#### S2.3.1 Removing shorter k-mers nested inside longer peak motifs

Cases in which both k-mer *x*[*s_i_, s_i_ + k*) (e.g., CATACTGAGA in Figure S1) and its sub-(k —m1 *—*m*2*)*-*mer *x*[*s_i_+m_1_, s_i_+k — m_2_*) (with m1 > 0, m2 > 0; e.g., TACTGAGA in Figure S1 with m1 = 2, m2 = 0) are individually identified can be resolved to report only the longer k-mer by removing any index *i* <E I (defined by Equation (S6)) if there exists *j* <E I such that the [kj]-mer interval starting at *s_j_* includes all of the [ki]-mer interval starting at si, thus retaining only:

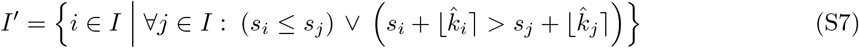

This can be done efficiently using an interval tree.

#### S2.3.2 Extending substring k-mers to match longer motifs from distinct peaks

Besides two cases of nested k-mers which may be removed from the reported motif set by the method of Section S2.3.1 (ATACTGAGA and TACTGAGA), Figure S1 also depicts a shorter k-mer ATACTG derived from a distinct occurrence of the same longer k-mer (CATACTGAGA). Because this distinct occurrence of the longer k-mer was not itself initially identified, the method of Section S2.3.1 does not remove the shorter substring k-mer from the motif set. However, such substring k-mers may be extended to the longer k-mer occurrence by the following method: for each *i* G I’, define the duplet

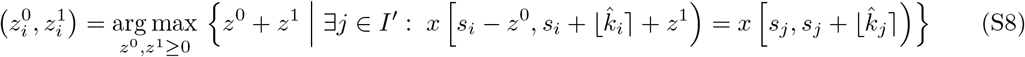

resolving any ties in the arg max in favor of maximal *z*^0^. Equation (S8) picks out the largest super-interval *s_i_* — *z*^0^, *s_i_* + [ki] + *z*^1^ *J* containing the interval *s_i_*, *s_i_* + [ki] *J* such that the extended 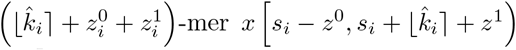 is equal to one of the already identified k-mers < *x s_j_*, *s_j_* + [kj~| *J j* G I’ >. (In the example of Figure S1, 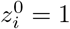 and 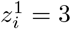, corresponding to the light gray highlighted characters surrounding the substring k-mer). Then

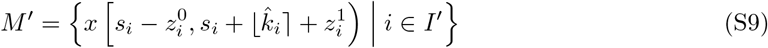

defines our motif set after removal of nested motifs.

### S2.4 Spatial smoothing to identify multi-motif domains

SArKS identifies candidate multi-motif domains (MMDs) through the application of a second round of kernel-smoothing over suffix positions *s_i_* within words:

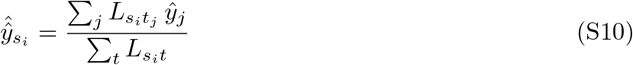

where we here use uniform kernels of the form

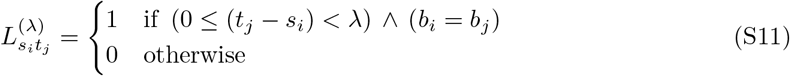

(generally with width λ ≠ *κ*) to search for regions of length λ with elevated densities of high-scoring motifs. Note that 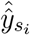 defined by Equation (S10) is indexed not by suffix array index *i* but by suffix array value *s_i_* giving the spatial position *s_i_* in the concatenated word *x*.

To use such spatial smoothing for motif selection/filtering, it is necessary to introduce a second threshold θ_spatial_, as the doubly-smoothed scores 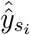 will generally be somewhat less dispersed than will be the singly-smoothed *ŷ_i_*. The threshold θ_spatial_ can be used to define an index set I_spatial_ similar to the manner in which I is defined by Equation (S6), but the task is more complex when we replace the single spatial position *s_i_* by a spatial window [*s_i_*, *s_i_* + λ).

Recalling the definitions of the negative/positive spatial shift operators η(*i*)*/ρ*(*i*) which yield the unique suffix array indexes corresponding to the spatial position immediately before/after si, so that *s_η_*(i) *=* s_i_ — 1 and *s_ρ_*(i) *=* s_i_ + 1, first define:

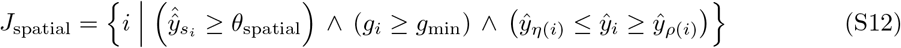

J_spatial_ represents the set of suffix array indices *i* corresponding to the left endpoints s_i_ of spatial windows [s_i_, s_i_ + λ) passing the filters for score threshold θ_spatial_ and minimum Gini impurity *g_min_*, and for which the sequence-smoothed score *yi* is at least as high as the spatially adjacent scores *ŷ_η_*(*i*) to the left and *ŷ_ρ_*(*i*) to the right.

Defining the left-directed distance *δj* from the suffix with sorted suffix array index *j* to the set Jspatial by

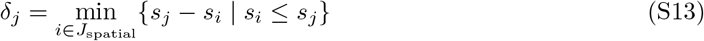

we define in turn the set of sorted suffix array indices Ispatial marking the starting positions of selected k-mers by:

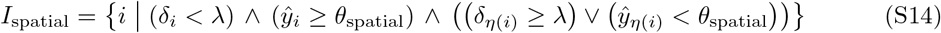

Equation (S14) identifies suffix array indices *i:* (1) whose spatial positions s_i_ fall within a spatial window [s_j_, *s_j_ +* λ) for some *j* G Jspatia_l_, (2) whose sequence-smoothed score *ŷ_i_* > θ_spatial_, and (3) for which the position s_i_ — 1 spatially to the left is either (3A) not in one of the spatial windows specified by Jspatial or (3B) has associated sequence-smoothed score *ŷ_η_*(i) *<* θspatial. This final criterion is included because we want to merge adjacent k-mers whose leftmost positions fall within the same selected spatial window.

This merging process is implemented by calculating for each index *i* G Ispatial the length

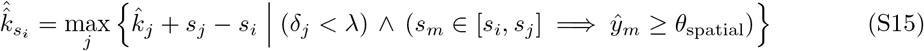

of the merged k-mer starting at i. Equation (S15) sets 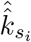 by selecting the right endpoint 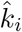 + *s_j_* of the k-mer beginning at *s_j_* to maximize the merged length 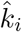 + *s_j_* — *s_i_* over all choices of *s_j_* for which every position sm between *s_i_* and *s_j_* has an acceptable sequence-smoothed score ym > θ_spatial_. It is then straightforward to obtain the motif set

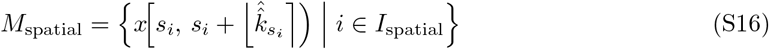

### S2.5 Permutation testing to establish significance of motif set

The significance of the observed correlation between the occurrence of the motifs uncovered by SArKS and the sequence scores *yb* can be evaluated by examining results obtained when the sequences *ω_b_* and the scores *yb* are independent of each other. To this end, the word scores *yb* are subjected to permutation π to define

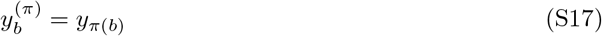

If the permutation π is randomly selected independently of both the sequences *ω_b_* and the scores *yb*, any true relationships between sequences and scores will be disrupted. This suggests a simple method for assessing the significance of motifs discovered using a given set of parameters (kernel half-width *κ*, θ, *g*_min_, etc.): generate *R* random permutations π*_r_* and for each permutation calculate scores 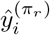 using Equation (4) (and also 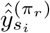 using Equation (S10) if spatial smoothing is employed) with *yb* replaced by *y*_π*b*_. In this manner one can estimate the distribution of scores under a null model in which there is no association between the sequences of the various words *ω_b_* and the scores *y_πb_*.

This method of significance testing also provides the motivation for the form of Equation (S4) in Section S2.2. Let Π be a random variable representing a random permutation and note that the random variables y_Π_(b) satisfy

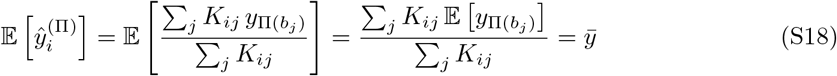

while, assuming that the number of words *n = |W|* is large enough that we may approximate y_Π_(*b*) J- y_Π_(*b’*) for *b = b’*,

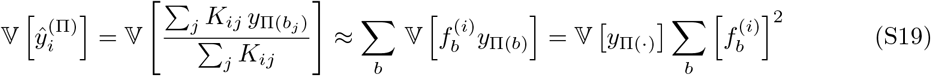

where *f*_b_^(i^ is defined by Equation (S3) and for all *b*

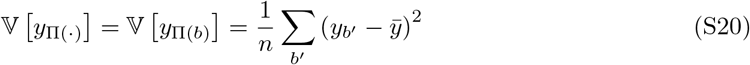

Equation (S19) then tells us that

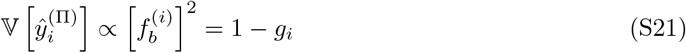

where the Gini impurity *g_i_* is defined by Equation (S4). Thus smaller values of *g_i_* imply higher variance 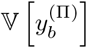 of the window-smoothed scores obtained under random permutation (with mean unchanged). This increased variance will lead to the requirement of larger cutoff values *θ* for reporting motifs discovered in the unpermuted data with a given degree of confidence unless positions *i* with *g_i_ < g*_min_ are filtered out as described in Section S2.2.

### S2.6 Permutation testing to set thresholds for multiple parameter combinations

Multiple combinations 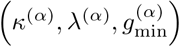 of the values of SArKS parameters may be explored (with α indexing the set of desired combinations); for example, Sections 3.2.1-3.2.2 discuss the parameters used in the benchmarking examples herein and the rationales for their selection. For any permutation π, let

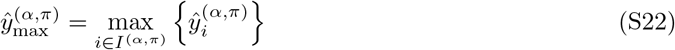

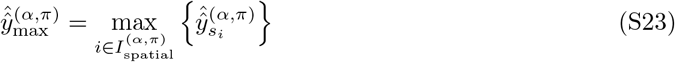

(i.e., 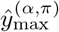 is the highest ltered sequence-smoothed score obtained ater permuting by π, while 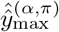 is similarly the highest filtered spatially-smoothed score). hen we suggest a simple method for setting thresholds θ(α) and 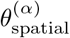 based on a set of randomly generated permutations {πr}:

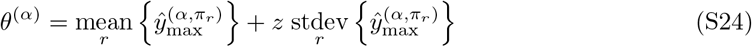

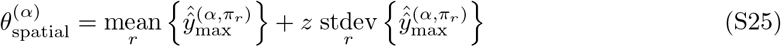

with higher values of z trading reduced sensitivity for lower false positive rates (in the examples analyzed in Section 3.2.1 we take z = 4). For the sake of simplicity we have generally used only one of these two thresholds for any particular combination of parameters α, setting θ(α) = —oo if κ(α) > 1 or 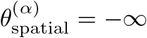 if *κ —* 1 (i.e., if spatial smoothing is not employed).

In order to characterize the false positive rate associated with the entire set of analyses across all of the parameter settings employed while controlling for multiple hypothesis testing, a family-wise error rate (FWER) e resulting from these thresholds can then be estimated by generating an independent set of *R’* permutations 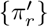 and counting the number of permutations 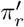 for which a nonempty set of k-mer motifs is identified using any of the parameter sets 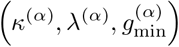. That is, writing

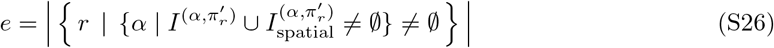

(where 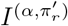 and 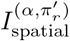 are defined respectively by Equation (S6) and Equation (S14) using the thresholds θ^(*α*^) and 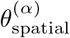 determined using the original permutation set {π_r_}) we can infer confidence intervals by noting that the random variable *E* instantiated in e satisfies *E* ~ Binom(R’, e) under the permutation test null hypothesis. We can thus derive confidence intervals (CIs) for the FWER (in the weak sense, as the permutation test represents a complete null hypothesis with no true positives (Farcomeni, 2008)) by applying the Clopper-Pearson method for estimation of binomial CIs.

### S2.7 RNA-seq expression analysis

#### S2.7.1 Assigning PV differential expression scores for Mo 2015 data set

In order to test SArKS, we selected the *M. musculus* neocortical neuron RNA-seq data set GSE63137

(Mo *et al.*, 2015) from Gene Expression Omnibus (GEO) database (Barrett *et al.*, 2013) (https://www.ncbi.nlm.nih.gov/geo/

This data set contains detailed transcriptomic and epigenetic information from three functionally and neurochemically distinct classes of pooled neocortical neurons: principal excitatory neurons, parvalbumin-positive (PV) GABAergic neurons, and vasoactive intestinal peptide-positive (VIP)

GABAergic neurons.

Because the position of the first exon can help pinpoint the TSS—and hence the DNA region containing the putative promoter—we reanalyzed the GSE63137 RNA-seq data using kallisto (Bray et *al*., 2015) to quantify and normalize transcript level expression against Ensembl mouse cDNA reference GRCm38. Transcript species were filtered by mean expression to focus on those for which reliable expression estimates could be made, retaining only transcripts for which at least 100 pseudocounts were obtained when summed across all samples and whose mean normalized expression met or exceeded the median of the transcript mean normalized expression levels. We also filtered out transcripts that showed low variance across the full sample set, retaining only those for which the estimated variance 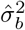 of normalized expression values met or exceeded median 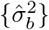 across all transcript species (Bourgon et *al*, 2010). In order to simplify downstream analysis, only the isoform with highest mean expression level across all samples was retained per gene. Finally, as based on chromatin accessibility data (Mo et *al*, 2015), only transcripts for which the transcription start sites were located within ATAC-seq peaks (i.e., were accessible) for all examined neuron classes were analyzed. This accessibility-based filter reduced the likelihood that epigenetic features, rather than regulatory sequences, determine the variations in gene expression between cell classes.

**Table S1.** Filters applied to select gene sets for SArKS analysis. The Mo 2015 data set (bulk RNA-seq) was realigned and analyzed at isoform level, hence counts in first column indicate distinct transcripts or isoforms. For the single-cell RNA-seq Close 2017 data set, the original gene-level alignment counts were analyzed; counts in second column indicate distinct genes. No variance filter was applied for the Close 2017 data set, as none of the 6,939 highly expressed genes exhibited low estimated variance. Epigenetic accessibility data was available for the Mo 2015 samples but not the Close 2017 samples.

|  | Mo 2015 | Close 2017 |
| --- | --- | --- |
| All | 111,669 | 23,045 |
| Detected | 73,912 | 15,490 |
| + Highly Expressed | 37,721 | 6,939 |
| + Highly Varying | 29,164 |  |
| - Duplicate Isoforms | 11,857 |  |
| + Accessible | 6,326 |  |
| Final SArKS Set | 6,326 | 6,939 |

Differential gene expression was assessed using normalized expression values via standard Student’s t-test (comparing data for PV neurons to data for excitatory and VIP neurons), with the resulting t-statistic providing an estimate of a gene’s enrichment in PV neurons (score *yb* for transcript *b*). One potential issue with the use of such *t*-statistics with small sample numbers—here, two samples associated with each neuron type—is that especially low within-group standard deviation estimates can result in very large magnitude *t*-statistics for a few genes. For example, the average estimated within-group standard deviation of the 76 genes with |*tb*| > 10 (with |*tb*| ranging up to a maximum value of 49.6) was less than 30% of the average within-group standard deviation of the full set of 6,326 analyzed genes (Table S1). Every one of the 76 genes with such high magnitude *t*-statistics had a within-group standard deviation estimate below the median value for the full gene set.

The phenomenon of low within-group variance estimates leading to inflated test statistics has previously led to the application of empirical Bayes methods (Smyth, 2004) using moderated *t*-statistics in place of standard *t*-statistics for calculating differential expression *p-*values. As we are here instead interested in using the *t*-statistics to derive word scores *yb*, for which no particular distributional assumptions are required, we have adopted a simpler approach to prevent the few very large magnitude *t*-statistics from unduly influencing motif discovery by applying a ceiling of 10 on the magnitude of *yb*:

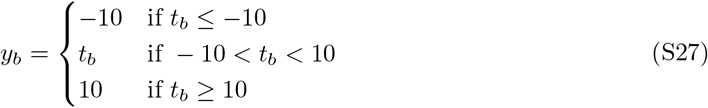

#### S2.7.2 Assigning DCX differential expression scores for Close 2017 data set

We also examined an RNA-seq data set comparing transcriptomes of in vitro-induced human embryonic stem cells and the resulting cultured interneurons (Close *et al.*, 2017). We applied SArKS to identify promoter motifs associated with elevated gene expression in doublecortin-positive (DCX+) interneurons. We restricted our analysis to the post-induction day 54 (D54) timepoint, where most of the DCX+ neurons were post-mitotic and GABAergic, and for which the largest total number of cells had been profiled, minimizing the within-group expression variations.

We used the normalized gene expression levels from GEO (Barrett *et al.*, 2013) records for this data set (accession GSE93593). We chose not to reanalyze the sequencing data in this case because we did not want to split the read counts per cell—which, given the large numbers of cells observed, tend to be much lower than read counts per sample in bulk RNA-seq—across multiple distinct transcripts for each gene. We found 15,490 genes for which (1) nonzero aligned read counts were detected in at least one (out of 585) analyzed cells and (2) a unique entry was found in the GRCh38 annotation of the human genome. We retained the 6,939 genes from this set for which the average aligned read count per cell was ≥ 25 for further analysis (Table S1). As the variance of the log2-transformed transcripts-per-million (TPM) normalized expression levels was quite high (≥ 1 for 6,852 of the 6,939 genes, ≥ 0.5 for all 6,939 genes), we did not apply any variance filter for this data set. As no epigenetic information was available, no accessibility filtering could be conducted.

For the filtered high-read-count gene set, differential expression was assessed via a simple two group t-test comparing the DCX+ cells to the DCX- cells and SArKS scores were assigned according to Equation (S27), just as was done for the Mo 2015 data set.

### S2.8 Specifications for running existing motif discovery algorithms

Existing algorithms were run at their default parameter settings (defined either within the source code or in associated documentation), with two exceptions: (1) MOTIF REGRESSOR was run searching only for motifs positively correlating with score to enable more direct comparison of its output with that of the other algorithms (none of which look for anticorrelated motifs by default). (2) STEME was run in discriminative mode using a high order Markov model on the negative sequences exactly as suggested in the online documentation (https://pythonhosted.org/STEME/using.html); however, STEME’s implementation requires pre-specification of the number of motifs to report, defaulting to a single motif if unspecified. Given that (Reid and Wernisch (2014)) extensively compared STEME to DREME with the finding that the two were generally comparable in performance, we took the upper bound on the observed DREME motif set size (10 motifs) as the number of motifs for STEME to report.

~~~
##
## DREME options: dreme \
-p $pos_seq_fasta \
-n $neg_seq_fasta \
-oc $out \
-png
##
## FIRE options: perl fire.pl \
--expfiles=$scores \
--exptype=continuous \
--fastafile_dna=$seq_fasta \
--seqlen_dna=$seq_len \
--nodups=1
##
## HOMER options: homer2 denovo \
-i $pos_seq_fasta \
-b $neg_seq_fasta
##
## MOTIFREGRESSOR options: MotifRegressor.pl \
$scores \
$seq_fasta \
null 1 1 2 1 0 50 250 50 250 5 15 50 30 \
$out ## _interpretation of MOTIFREGRESSOR parameters above_
## null : background sequence distribution to be computed based on input sequences ## 1 : use column 1 from $scores to rank sequences ## 1 : use column 1 from $scores to perform regression ## 2 : data does need to be further log-transformed
## 1 : look for motifs in high-scoring (as opposed to low-scoring) sequences ## 0 : select fixed count of top motifs (as opposed to setting fixed score threshold) ## 50* : number of initial top motifs
## 250* : number of sequences with high values for confirmation ## 50* : (ignored since we are only interested in high-scoring motifs) ## 250* : (ignored since we are only interested in high-scoring motifs) ## 5* : minimum motif width ## 15* : maximum motif width ## 50* : number of seed candidate motifs ## 30* : number of motifs reported before regression
## _all parameters marked with * were set at the example (e.g.) values given in ## MOTIFREGRESSOR’s README.MR file_
##
## STEME options: steme \
--output-dir=$out \
--num-motifs=10 \
--bg-model-order=5 \
--bg-fastafile=$neg_seq_fasta \
$pos_seq_fasta ## _see https://pythonhosted.org/STEME/using.html_
~~~

**Table S2.** Unpermuted scores consistently exceed permuted scores only when motif is present. Distribution of simulated differences *ŷ_max_* – *ŷ*^(π)^ obtained by suffix array kernel smoothing when CATACTGAGA motif is embedded into 10 high score sequences (motif (+) column) or when it is not (motif (-) column). The values *ŷ_max_* were calculated by smoothing the unpermuted sequence scores *yb*, while the values 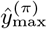 were obtained using permuted sequence scores y_π(b)_. When motif is included, *ŷ_max_* – *ŷ*^(π)^ tends to be positive—i.e., unpermuted smoothed scores usually exceed permuted—while when motifs are not present the distribution is symmetric about 0, reflecting the lack of signal for SArKS to detect.

| $\hat{y}_{\max} - \hat{y}_{\max}^{(\pi)}$ | motif (+) | motif (-) |
| --- | --- | --- |
| -29 | 0 | 7 |
| -19 | 0 | 275 |
| 0 | 29 | 479 |
| 19 | 551 | 229 |
| 29 | 420 | 10 |

## S3 Results and Discussion

### S3.1 Illustration of SArKS using simulated data

To demonstrate that the results of Section 3.1 are not a quirk of a single simulation, we repeated the process of (1) generating 30 random sequences, embedding the motif CATACTGAGA into the last 10 sequences, and (2) applying SArKS to the sequences and sequence scores 1000 times. In 971 iterations, the maximum value

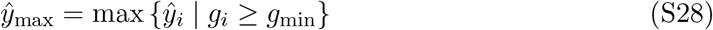

calculated using the unpermuted sequence scores exceeded the maximum value

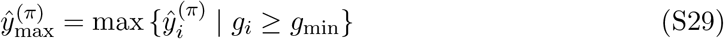

obtained using one set of randomly permuted sequence scores per iteration. The full distribution of the differences 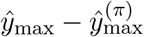 is shown in the motif (+) column of Table S2.

We also examined the results of SArKS applied to simulated data in which no motif was present to find; for this purpose, we repeated an amended version of the simulation process 1000 times, omitting the motif embeddings. The distribution of 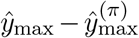 for these no-motif simulations is presented in the motif (-) column of Table S2. In this case, *ŷ_max_* exceeded 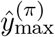 in only 239 of the simulations, while 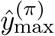 exceeded *ŷ_max_* in 282 simulations, with equality between the two holding in the remaining 479 iterations. The symmetry of the distribution of 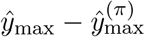 around 0 in the motif (-) case is to be expected since the scores *yb* are independent of the sequences *ω_b_* whether permuted or not if no motifs are included. By contrast, the strong asymmetry of the distribution of 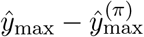 when the motif is present demonstrates the ability of the permutation approach to differentiate a true signal from background noise.

**Figure S2.**
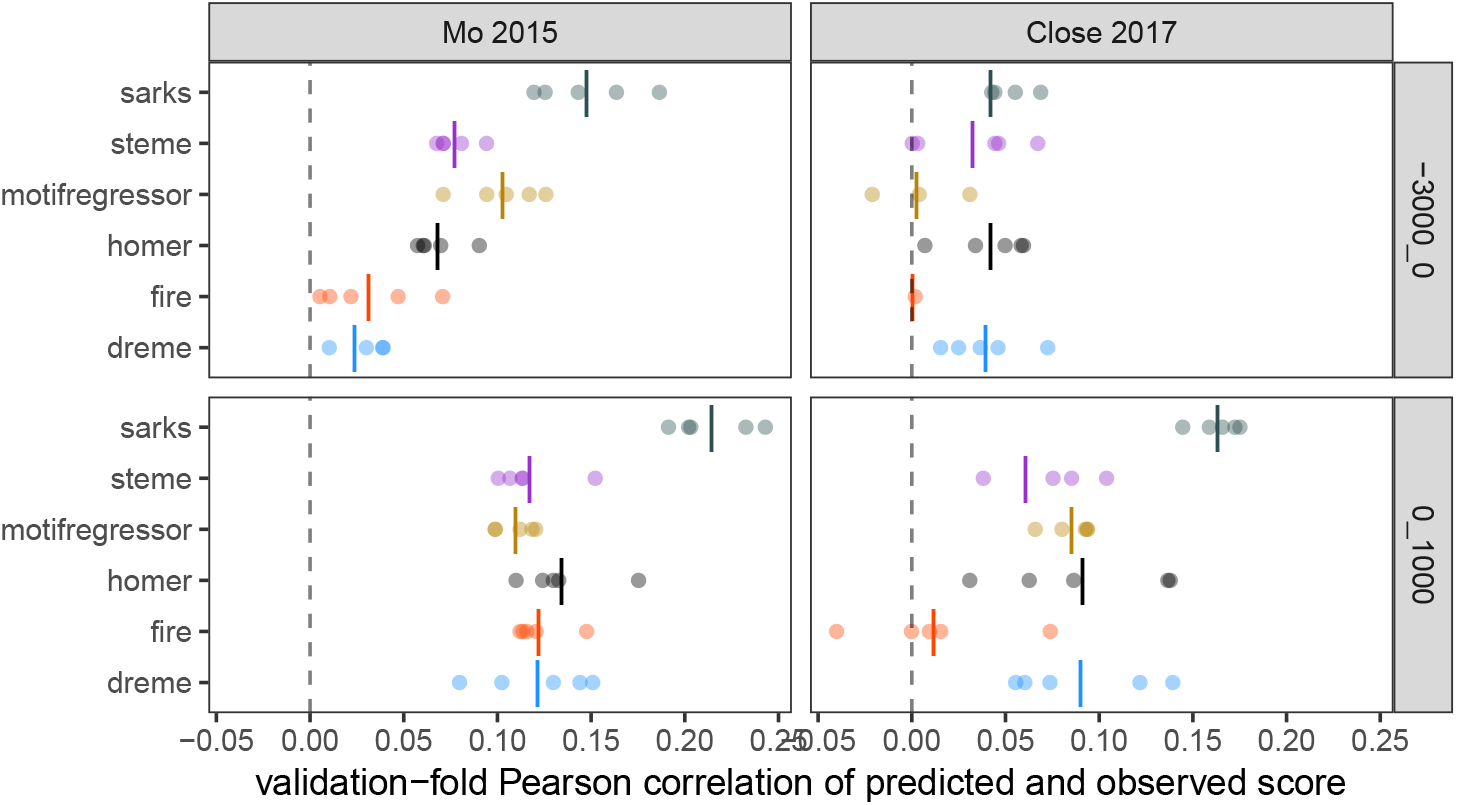
Motif regression model predictions correlate with gene specificity scores in held-out validation subsamples. Each of five cross-validation folds is plotted as separate point for each algorithm; fewer than five points are shown for corresponding algorithm if it failed to identify any motifs in one or more cross-validation folds. Upper panels: regions 3kb upstream of TSS; lower panels: regions 1kb downstream of TSS. Vertical lines indicate mean Pearson correlation across all folds (including a value of 0 for any fold in which algorithm failed to identify any motifs).

### S3.2 Uncovering promoter motifs associated with differential gene expression

#### S3.2.1 Benchmark comparisons to existing algorithms

The cross-validated regression modeling strategy described in Section 3.2.3 builds a single regression model based on the concatenated upstream and downstream motif count vectors. We also built two more separate regression models—one using as feature set only the upstream motif counts, the other only the downstream motif counts—for each of the two data sets, obtaining the results presented in Figure S2. SArKS generally outperformed the other algorithms in these comparisons, though for the upstream analysis of the Close 2017 data, DREME and HOMER offer similar performance; all of the algorithms have their poorest performance in this particular analysis. For both data sets the regression model predictions on the held-out validation folds are noticeably better in the downstream analyses than the upstream analyses, as discussed in Section 3.2.3.

We used `tomtom` (Gupta *et al.*, 2007) to compare the pooled motif sets identified by each algorithm and detected overlap between motifs sets by algorithm (S3). There exists a significantly similar (q ≤ 0.1) SArKS-identified motif for the majority of motifs identified by any of the existing algorithms in the Mo 2015 data set. For the Close 2017 data set, at least 50% of the motifs identified by DREME, FIRE, MOTIF REGRESSOR, and STEME can be paired with a significantly similar SArKS motif, though this is true for only 39% of HOMER-identified motifs.

An alternative benchmarking approach is to compare the motifs identified algorithmically to databases of known TF-binding motifs, such as JASPAR (Mathelier *et al.*, 2015). For the presence of a TF-binding site to be biologically relevant in a cell, it is necessary for the TF itself to be present as well. In the context of our analysis of the two RNA-seq data sets, we checked whether or not mRNA encoding TFs whose binding sites were similar to discovered motifs are enriched among either the PV neuron (Mo 2015) or DCX+ cell (Close 2017) transcripts. We classified a TF gene as enriched if there was at least one distinct mRNA transcript for the gene with (1) at least 100 reads (or pseudocounts for the Mo 2015 set) and (2) for which the mean TPM-normalized estimated expression level in either the PV samples (Mo 2015) or DCX+ cells (Close 2017) ≥ the median of the genewise means for all measured transcripts/genes in the relevant data set. Figure S4A is similar to a receiver-operating characteristic plot in which the motif discovery algorithms are regarded as classifying JASPAR motifs as positive when they show sufficient similarity to any of the discovered motifs; the distance of a point above the diagonal indicates the degree to which an algorithm preferentially identifies binding motifs for TFs showing high RNA-seq expression levels in the target cell population. SArKS identifies motifs similar to a larger fraction of JASPAR than do the other algorithms while maintaining a preference for motifs for highly expressed TFs.

**Figure S3.**
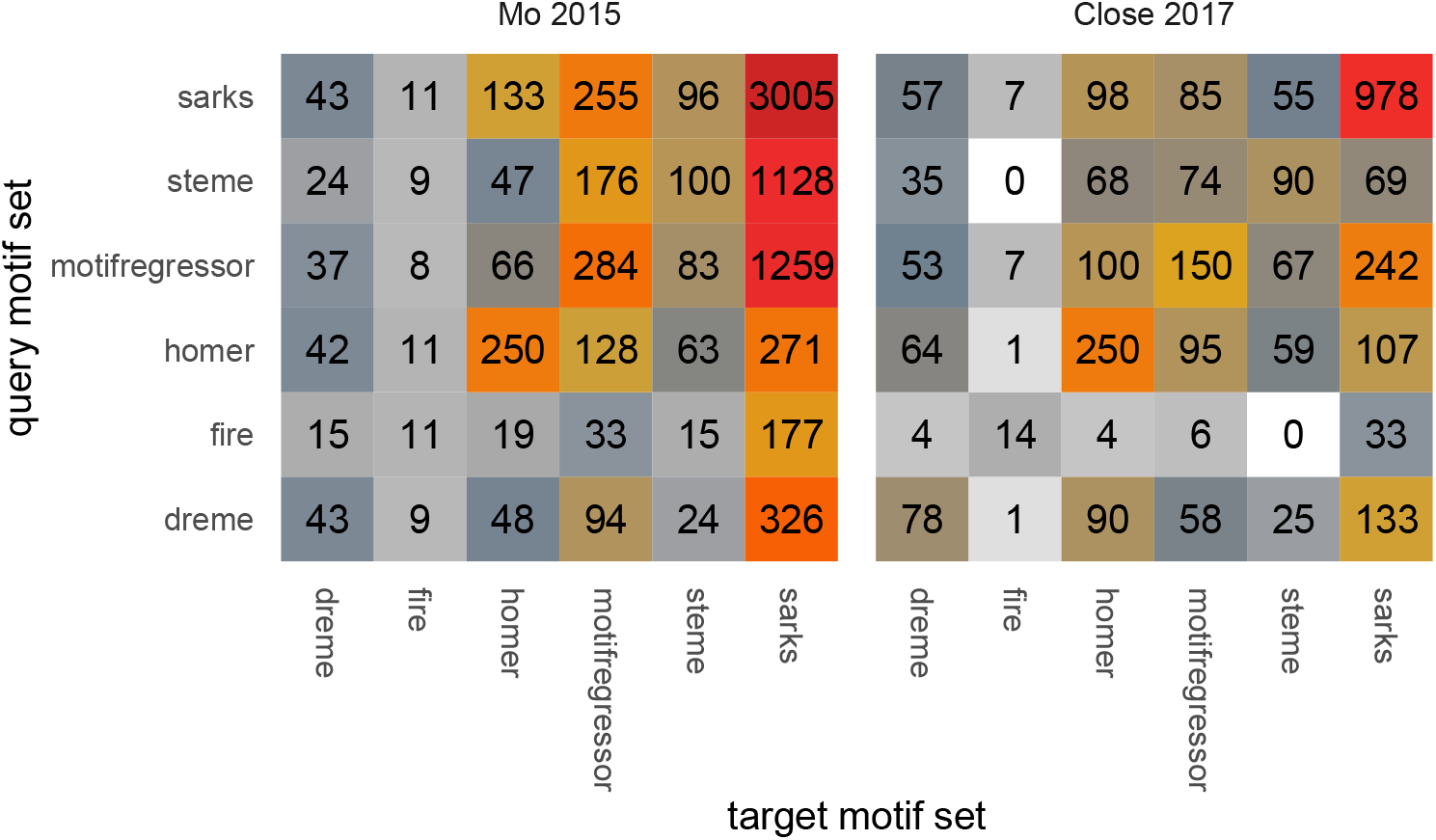
Counting motif similarities shows substantial overlap between algorithms. Each cell indicates the count of motifs identified by the target motif set algorithm for which there is a motif in the query set with significant `tomtom` similarity (q ≤ 0.1). Cells are colored according to the numbers they contain.

Figure S4B illustrates the overlaps between the sets of JASPAR motifs with similarities among the motifs identified by the motif discovery algorithms: For all algorithms applied to the Close 2017 data set and all but HOMER in the Mo 2015 data set, the set of JASPAR motifs with significant similarity (q ≤ 0.1) to one of the algorithm-identified motifs overlaps by more than 50% with the set of JASPAR motifs significantly similar to a SArKS motif. The degree of overlap between the JASPAR matches among the various algorithm motif sets tends to be higher than the degree of overlap directly between the motif sets themselves. This suggests that the presence of a similar JASPAR motif may provide supporting evidence that a given detected motif is a not a false positive.

**Figure S4.**
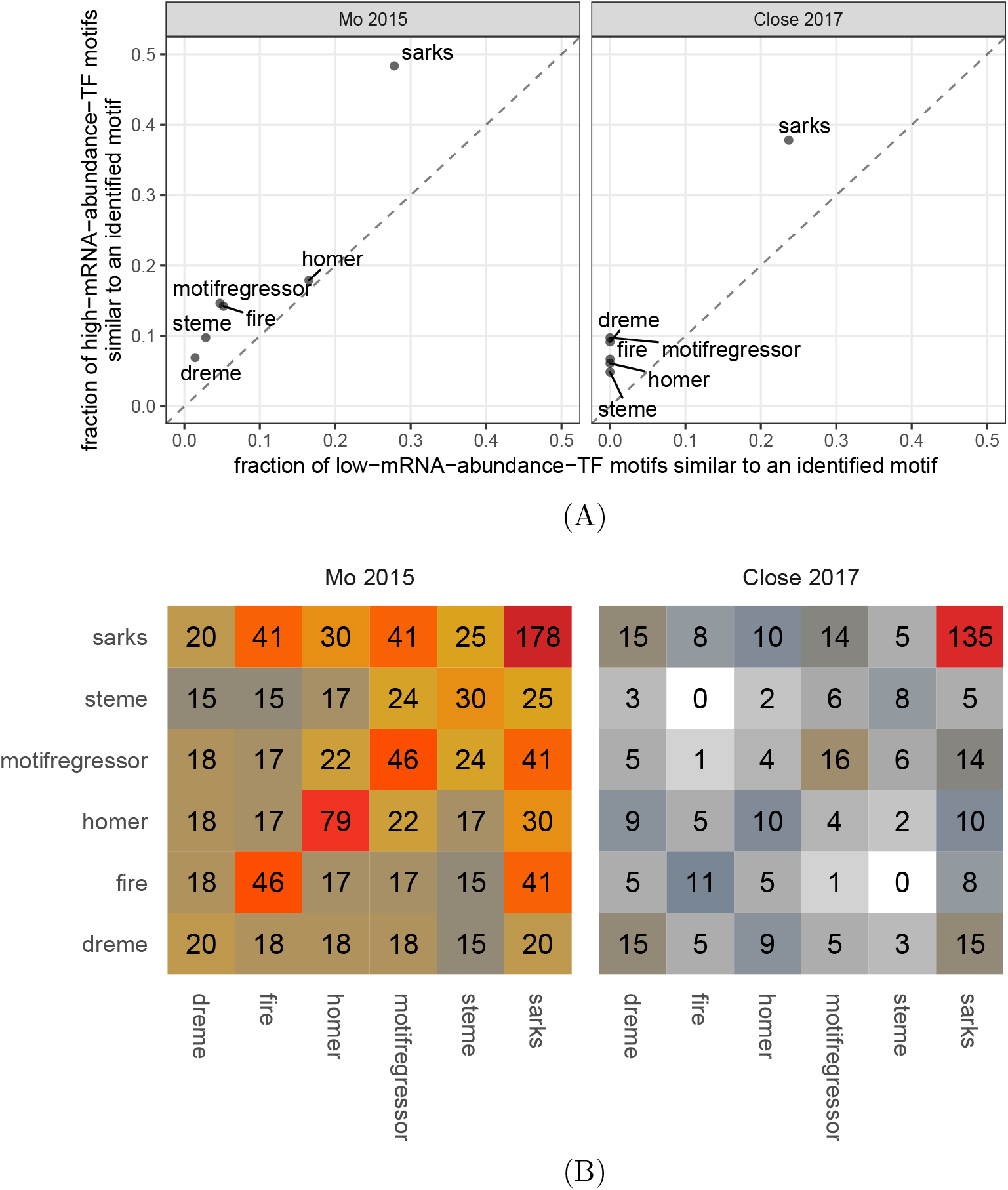
Discovered motifs overlap with known transcription factor binding sites. **(A)** Fractions of JASPAR-annotated TFs for which the algorithms indicated identified a motif with significant `tomtom` similarity (q ≤ 0.1) to the corresponding JASPAR binding motif. Vertical axis: fractions calculated using only the JASPAR-annotated TFs whose measured expression in either PV neurons (left panel) or DCX+ cells (right panel) were in top 50% by mean normalized expression (TPM) and had at least 100 associated reads. Horizontal axis: fractions calculated using only the remaining JASPAR-annotated TFs with measured expression below these expression filters. **(B)** Each cell indicates the count of JASPAR motifs for which there is a motif in both of the indicated algorithm motif sets with significant `tomtom` similarity (q ≤ 0.1). Cells are colored according to the numbers they contain.

**Table S3.**
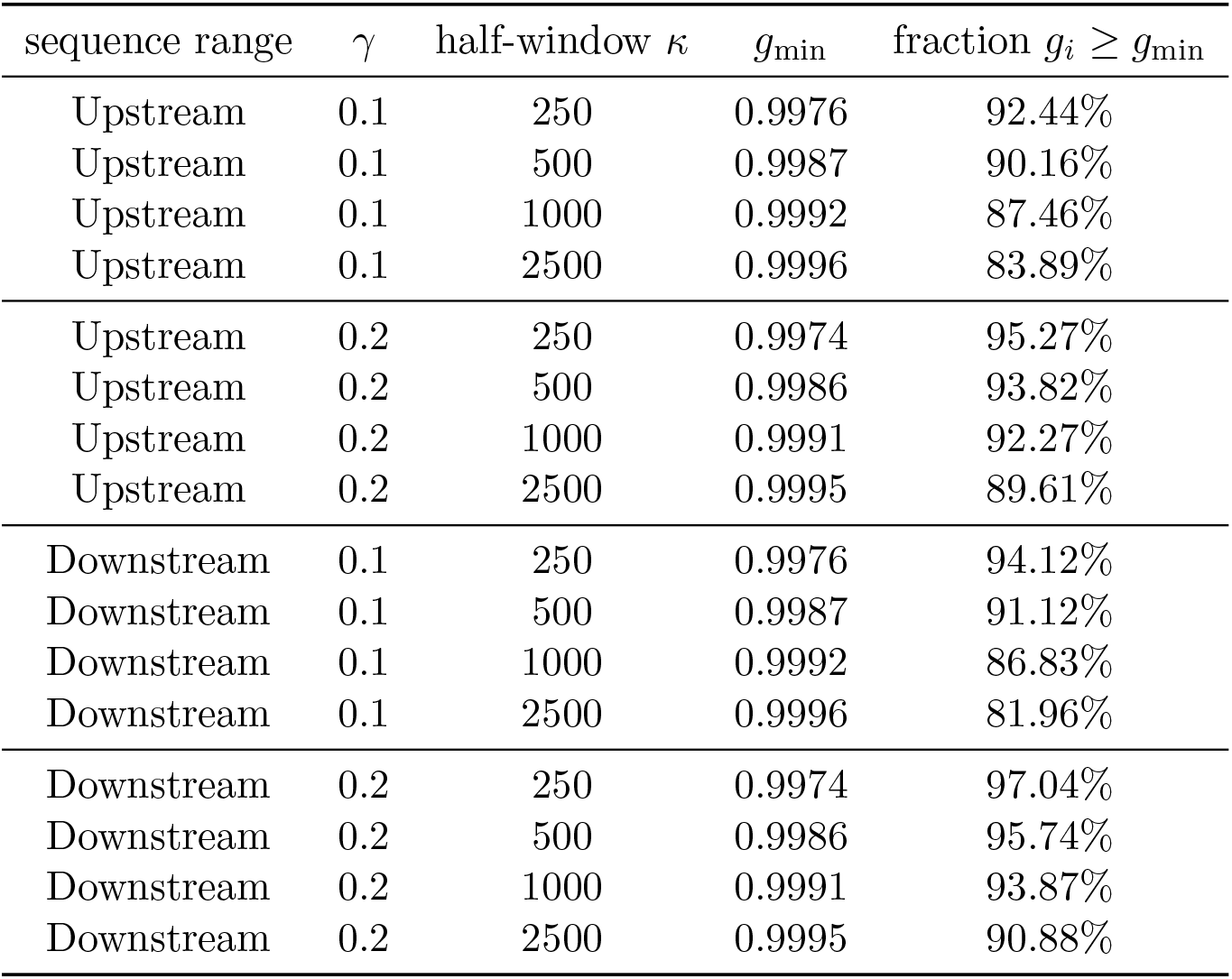
Gini index filters remove small fractions of suffix array positions. Fraction of suffix array positions *i* for which Gini impurity values *g_i_* ≥ *g_min_*, with *g_min_* selected according to Equation (S5) (applied to Mo 2015 data set).

| sequence range | $\gamma$ | half-window $\kappa$ | $g_{\min}$ | fraction $g_i \geq g_{\min}$ |
| --- | --- | --- | --- | --- |
| Upstream | 0.1 | 250 | 0.9976 | 92.44% |
| Upstream | 0.1 | 500 | 0.9987 | 90.16% |
| Upstream | 0.1 | 1000 | 0.9992 | 87.46% |
| Upstream | 0.1 | 2500 | 0.9996 | 83.89% |
| Upstream | 0.2 | 250 | 0.9974 | 95.27% |
| Upstream | 0.2 | 500 | 0.9986 | 93.82% |
| Upstream | 0.2 | 1000 | 0.9991 | 92.27% |
| Upstream | 0.2 | 2500 | 0.9995 | 89.61% |
| Downstream | 0.1 | 250 | 0.9976 | 94.12% |
| Downstream | 0.1 | 500 | 0.9987 | 91.12% |
| Downstream | 0.1 | 1000 | 0.9992 | 86.83% |
| Downstream | 0.1 | 2500 | 0.9996 | 81.96% |
| Downstream | 0.2 | 250 | 0.9974 | 97.04% |
| Downstream | 0.2 | 500 | 0.9986 | 95.74% |
| Downstream | 0.2 | 1000 | 0.9991 | 93.87% |
| Downstream | 0.2 | 2500 | 0.9995 | 90.88% |

#### S3.2.2 Case study: analysis of SArKS results for Mo 2015 data set

The values of *g_min_* obtained for the analysis of the Mo 2015 gene set (6,326 genes remaining after application of filters described in Section S2.7.1), along with the fraction of suffix array index values *i* for which *g_i_* ≥ *g_min_*, are listed in Table S3.

##### S3.2.2.1 Top motif identified in sequences downstream of TSS

The highest *ŷi* value obtained—detected in the downstream sequence analysis using *κ* = 250, λ = 0, and 7 = 1.1 in the downstream region analysis—corresponded to the fc-mer TGACCTTG. This fc-mer is very similar to a number of JASPAR TF-binding motifs. The strongest matches are to the binding motifs of ESRRB (*q* = 0.00078), ESRRA (*q* = 0.00078), and ESRRG (*q* = 0.00301). In fact a large fraction of the motifs associated with identified peaks in *ŷ_i_* identified in the downstream analysis exhibit significant similarity to one of the JASPAR motifs ESRRB, ESRRA, or ESRRG, as is illustrated in Figure S5B. The ESRR(A/B/G) TFs are all members of the estrogen-related receptor family; there is evidence that these receptors are involved in brain functions including synaptic transmission, neuronal firing, and mitochondrial biogenesis (Saito and Cui, 2018). This particular set of motifs may also help to explain the overall stronger performance of all of the motif discovery algorithms using the downstream sequences relative to the upstream sequences (Figure 3A), as we noted that all of these algorithms identified motifs similar to each of these JASPAR motifs (Section 3.2.3).

**Figure S5.**
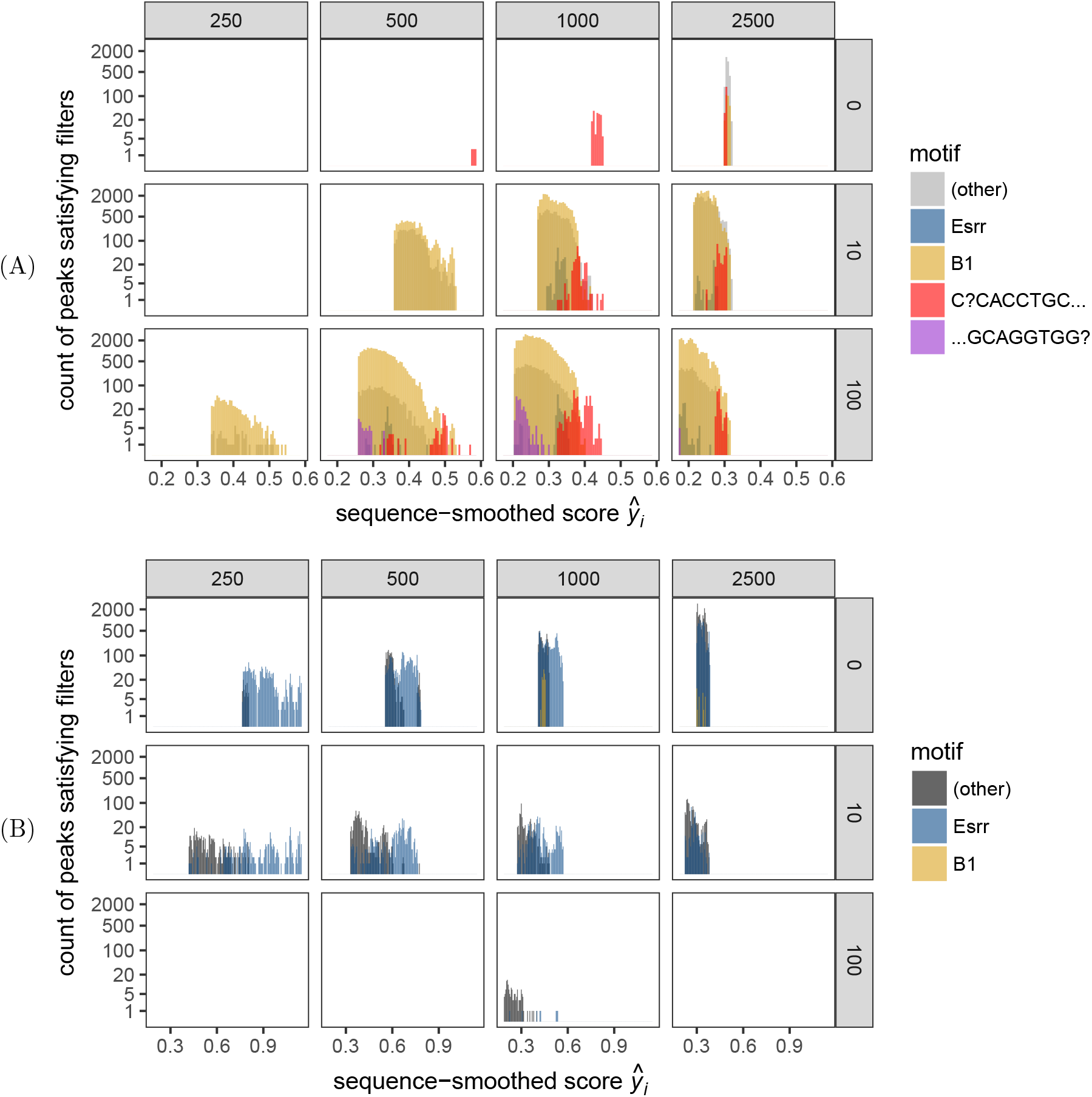
Contributions of top motifs to peak composition. **(A)** Log-scaled histograms of peaks *i ∊ I* (or I_spatial_ when spatial smoothing is employed) identified in **upstream** analysis for which corresponding k-mer motifs: (1) are prefixed with CACCTGC or CCACCTGC (indicated in red) or are suffixed by the reverse complement sequences GCAGGTG or GCAGGTGG (purple); (2) are otherwise spatially located within a `blast` hit to the B1 SINE sequence (gold); (3) exhibited significant `tomtom` similarity (*q* ≤ 0.1) to one of the JASPAR motifs ESRRA, ESRRB, or ESRRG (blue); or (4) did not satisfy any of the above criteria (gray). Horizontal panels: half-window κ values used in analysis; vertical panels: spatial smoothing length λ. (B) Log-scaled histograms of peaks identified in downstream analysis; color coding is as in (A) except that black replaces gray. C?CACCTGC and its reverse complement do not occur in downstream peak set.

**Figure S6.**
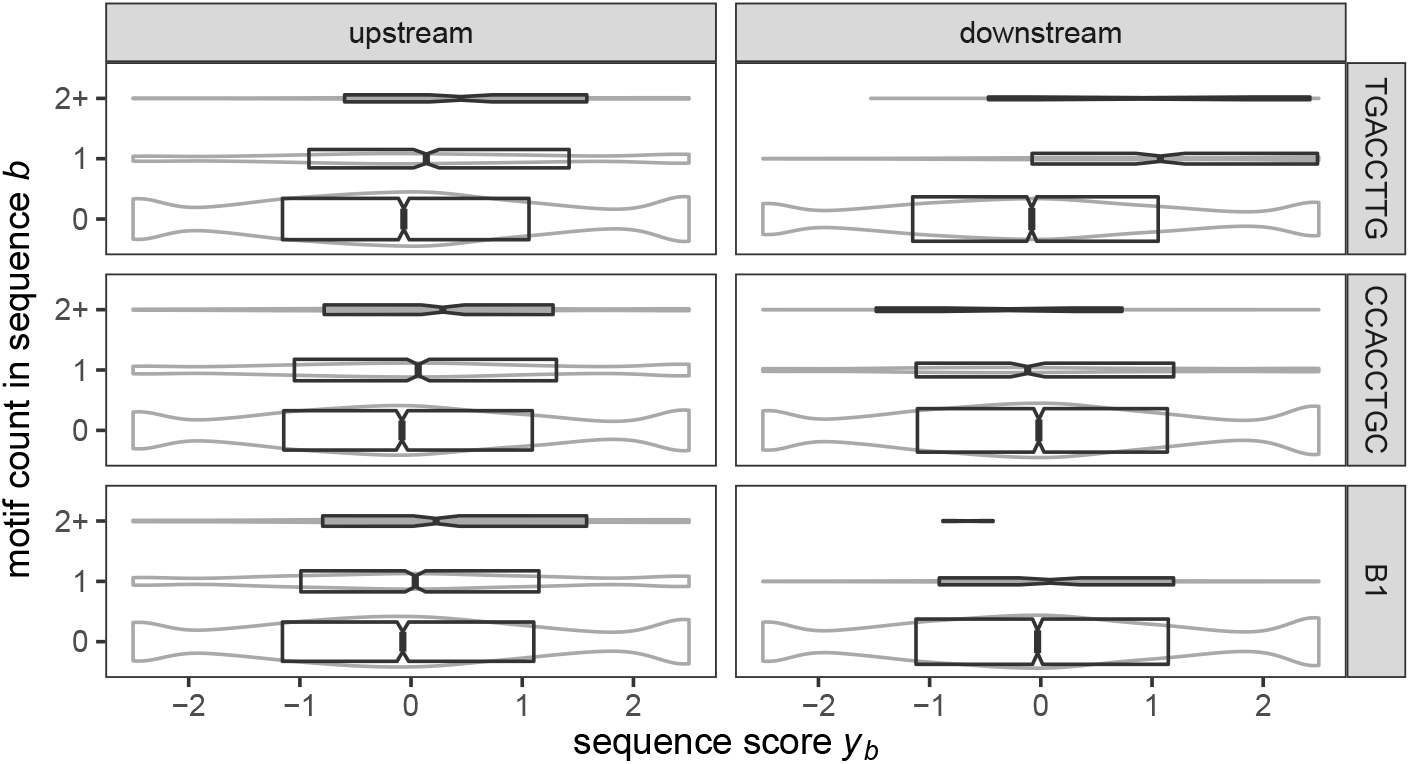
Sequence score *yb* distributions are shifted upwards in sequences containing one or more copies of top SArKS motifs. Each plot shows distribution of sequence scores split by the number of occurrences of motif indicated by the row label (TGACCTTG, CCACCTGC, or B1) found in the sequence range indicated by the column label (upstream or downstream). The first two motifs—k-mers TGACCTTG and CCACCTGC—were counted using regular expression matching (allowing matches on either forward or reverse strands), while B1 counts were assessed using `blastn` (percent identity > 90%, alignment length ≥ 70). Distributions are summarized by notched boxplots (area scaled to square root of sequence count; notch width is 1.57 times the interquartile range (IQR) divided by square root of sequence count) laid over kernel density estimates drawn as gray violins (area scaled to sequence count). Scores < — 2.5 or > 2.5 are lumped together into lower and upper ends of distribution, respectively. The score distribution for both TGACCTTG-positive upstream sequences and TGACCTTG-positive downstream sequences is shifted upwards, though the shift is notably larger in the downstream sequences. For the top motif CCACCTGC derived from analysis of the upstream sequences, however, the scores for the downstream sequences containing the k-mers do not show the same upward shift in the score distribution.

##### S3.2.2.2 Top motifs identified in sequences upstream of TSS

ESRRB/ESRRA/ESRRG binding motifs were also identified by SArKS analysis of the upstream sequences, but they did not account for either the highest scores *ŷ_i_* nor did they correspond to a large fraction of the overall k-mer motif sets discovered (Figure S5A). Figure S6 sheds some light on this: the distribution of sequence scores *yb* for downstream sequences containing one or more copies of the top SArKS octamer TGACCTTG is shifted upward to a much higher degree than is the the distribution of *yb* values for upstream sequences containing TGACCTTG.

Instead, For five of the 12 distinct combinations of smoothing half-window *κ* and spatial window λ investigated using SArKS, the k-mer CCACCTGC was identified at the positions *s_i_* with maximal values of *ŷ_i_* (the k-mers GCACACCTT, TGGAACTCACT, CCTGGAAC, and CAGCCTGG (identified using two distinct parameter combinations at the same suffix index *i*) were associated the highest *ŷ_i_* values using the remaining seven parameter combinations). The octamer CCACCTGC contains the canonical core recognition E-box sequence CANNTG (specifically, the E12-box variant CACCTG (Bouard *et al.,*, 2016); we note that the significant SArKS peak set contains many peaks corresponding to the 7-mer CACCTGC as well as the longer octamer adding the extra initial C). Comparison of CCACCTGC with known motifs from the JASPAR database using `tomtom` finds some evidence of similarity to 10

TF-binding motifs (SNAI2, MAX, SCRT2, SCRT1, TCF3, MNT, Id2, MAX::MYC, TCF4, and FIGLA; q-values of 0.14 for each), though no similarities significant at *q* ≤ 0.1. Unlike the case for the ESRR(A/B/G) motifs discovered in the downstream analyses, for which all of the benchmarked algorithms detected a matching motif, only one of the other algorithms (HOMER) detected a motif similar to either CACCTGC or CCACCTGC (`tomtom` *q* ≤ 0.1; no other algorithm produced any motifs matching even at *q* ≤ 0.5).

##### S3.2.2.3 B1 SINE sequence identified through MMD analysis

As the octamer CCACCTGC was identified in analyses with spatial window length λ ranging up to 100, we performed a multiple sequence alignment using muscle (Edgar, 2004) of the 100-mers *x*[*s_i_* — 50, *s_i_* + 50) for these positions *s_i_* (Figure S7A); three of the five 100-mers thus aligned were very similar (Levenshtein distance ≤ 7) to the 99-mer consensus sequence constructed. Furthermore, the consensus sequence also contains CCTGGAAC and CCAGGCTG (reverse complement of CAGCCTGG).

A `blast` screen of known repeated elements in the mouse genome for a consensus sequence uncovered a 93.9% identical base pair stretch of the B1 short interspersed element (SINE) sequence (SINEBase (Vassetzky and Kramerov, 2013)). The B1 SINE family consists of retrotransposon-derived sequences appearing throughout the mouse genome, especially upstream and within introns of genes implicated in DNA remodeling and expression regulation (Tsirigos and Rigoutsos, 2009). Additional observations have further suggested that SINEs function as transcriptional enhancers (Ichiyanagi, 2013; Elbarbary et al, 2016; Ge, 2017).

Figure S5A indicates the number of SArKS-identified peaks that fall within `blast` hits between the upstream sequences *ω_b_* and the B1 SINE consensus sequence as well as the numbers of peaks corresponding to the top motifs discussed above. The upstream SArKS peaks derived from analyses involving spatial-smoothing (λ ∊ {10,100}) are dominated by B1 sequences, many including the CCACCTGC motif or its reverse-complement.

##### S3.2.2.4 SArKS motifs correspond to variations on B1 sequence

Figure S7B provides a more detailed view of these peak counts by splitting them out by position to which the corresponding k-mers align to the B1 consensus and by whether they are matched or mismatched to the B1 consensus at each position. The k-mer CCACCTGC itself is not quite a perfect match to the canonical B1, containing a single base substitution away from the octamer CCGCCTGC whose reverse complement GCAGGCGG is found at positions 49-56 of the SINEBase B1 sequence. This substitution is responsible for the peak at position 54 in the mismatch counts in Figure S7B—one of the few positions at which there are more mismatches than matches. This G to A substitution creates the above noted E-box sequence CANNTG, while the unmodified octamer CCGCCTGC does not match any JASPAR motifs at *q* ≤ 0.5. This highlights the ability of SArKS to discover potentially functionally significant variations within a recurring sequence.

One of the remaining top upstream k-mer motifs mentioned above, GCACACCTT, similarly matches the nonamer GCACGCCTT spanning positions 15-23 of the SINEBase B1 sequence, but with a single G to A substitution. The modified nonamer GCACACCTT identified by SArKS shows significant similarity (tied *q* values of 0.038) to several JASPAR motifs (TBX21, EOMES, TBX15, TBX1, and TBX2), while the unmodified B1 nonamer GCACGCCTT again shows no similarity to any JASPAR motifs at *q* ≤ 0.5, again suggestive that specific SINE variants may promote differential gene expression.

**Figure S7.**
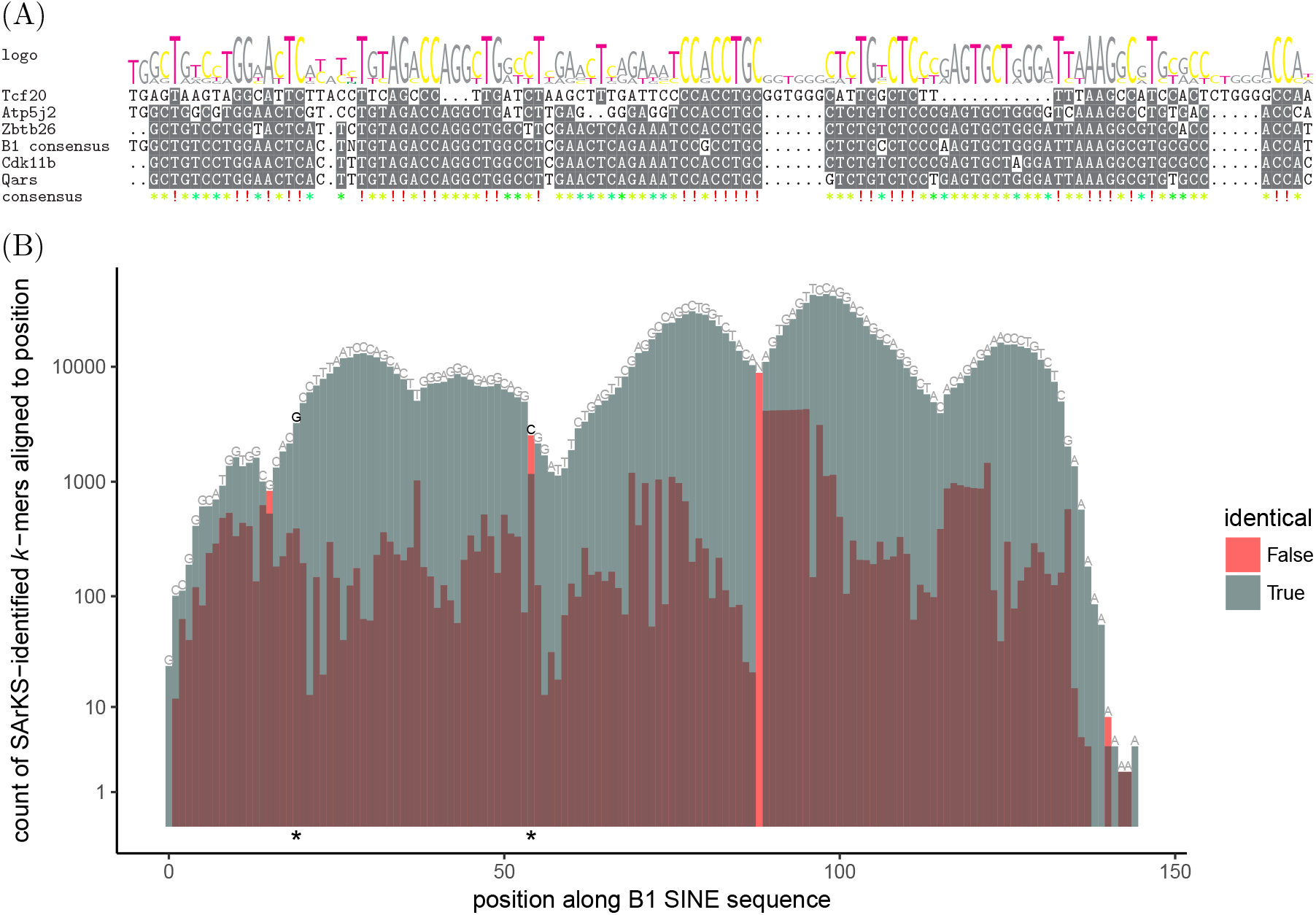
SArKS-discovered motifs within B1 SINE elements. **(A)** Multiple sequence alignment (muscle) of 100-mers surrounding top CCACCTGC motif peaks with reverse-complement of B1 consensus sequence. Associated genes are indicated to the left. Gray highlighting: ≥ 50% agreement in the multiple sequence alignment. **(B)** Number of upstream motif k-mer peaks in B1 regions that align to each position within the B1 sequence. Gray bars: number of peak k-mers derived from upstream sequence regions for which a `blast` hit (percent identity ≥ 90%, alignment length ≥ 70) to B1 was found and for which an alignment of the k-mer to B1 aligned a matching base at the position in question. Red bars: number of k-mers within B1 `blast` hits which align against B1 with a mismatched base at the position in question. Above each bar is a label indicating the B1 consensus base at that position. Note that the lack of a gray bar at position 89 results from the lack of consensus base for B1 at this position (marked by N above the red bar), so that all k-mers that align against this position must produce a mismatch. The consensus base labels are drawn darker and the bars are marked with an asterisk at positions (19 and 54) where two of the top SArKS peaks exhibit changes compared to the B1 consensus sequence. While essentially the entirety of the B1 consensus is represented by identified k-mer motifs, there is more variation away from the consensus towards the left end and at a couple of isolated positions further in than along most of the length of B1.

**Table S4.** SArKS-based regression modeling assists in selecting candidate upstream regions for promoting PV-specific expression. Number of distinct transcripts remaining after sequential application of described filters. Annotation of transcripts as protein coding or otherwise taken from Ensembl GRCm38 (Aken *et al.*, 2016). Expression levels were considered high in PV samples if the average within-PV value of log2(TPM + 1) ≥ log2(10 + 1), while expression levels were considered low in non-PV samples if the average non-PV log2(TPM + 1) < log2(10 + 1). Log-ratios were calculated as the difference of the PV-averaged- and non-PV-averaged-log2(TPM + 1) values, so that a log-ratio of one represents at least a two-fold increase in expression levels. SArKS regression scores were calculated using a ridge regression model built using counts of all k-mer motifs identified by SArKS applied to 3kb upstream promoter regions.

| Distinct Transcripts | Count |
| --- | --- |
| All analyzed | 6,326 |
| + Protein coding | 5,017 |
| + High expression in PV | 1,595 |
| + Low expression in non-PV | 196 |
| + PV : non-PV log-ratio $\geq 1$ | 92 |
| + Top 5% SArKS regression score | 13 |
| + Top 5% $t$-statistic score $y_b$ | 11 |

##### S3.2.2.5 SArKS-based candidate promoter selection

Finally, to illustrate how SArKS can be used to help select candidate regulatory regions for promoting specific expression patterns, we again constructed a ridge regression model based on the counts of SArKS-identified k-mer motifs. We applied the same modeling strategy as described in Section 3.2.3 to the promoter regions defined by the 3,000 base pairs immediately preceding the TSSs of each of the 6,326 distinct analyzed transcripts. Each distinct transcript was then assigned a score by resubstitution into the resulting regression fit. Table S4 shows a sequence of filters in which these regression scores were applied alongside other relevant criteria to select candidate PV-specific promoter regions. The promoter regions associated with the genes ATP5SL, GPRC5B, IFT27, KCNH2, MAFB, PAQR4, SLC29A2, SYT2, TBC1D2B, TMEM186, and TTC39A comprise the 11 candidates (from the final row of Table S4) selected for further experimental validation. Table S5 shows which of the top motifs discussed above are present in each of the candidate promoter regions: all regions except those for GPRC5B and MAFB contain at least one match for the ESRRB motif, while several also contain one or more copies of the E-box sequence and/or a match to the B1 SINE sequence. The promoter for IFT27 contains a match to a variant the B1 sequence with the substitution creating the E-box sequence CACCTGC. It is worth noting that there are many other SArKS motifs contributing to the promoter ranking model used here. Indeed, in accord with the principle that there is likely to be more than one way for combinations of motifs to achieve expression specificity, the candidate promoters for the genes GPRC5B and MAFB are ranked highly based exclusively on motifs other than the highest scoring ones.

**Table S5.**
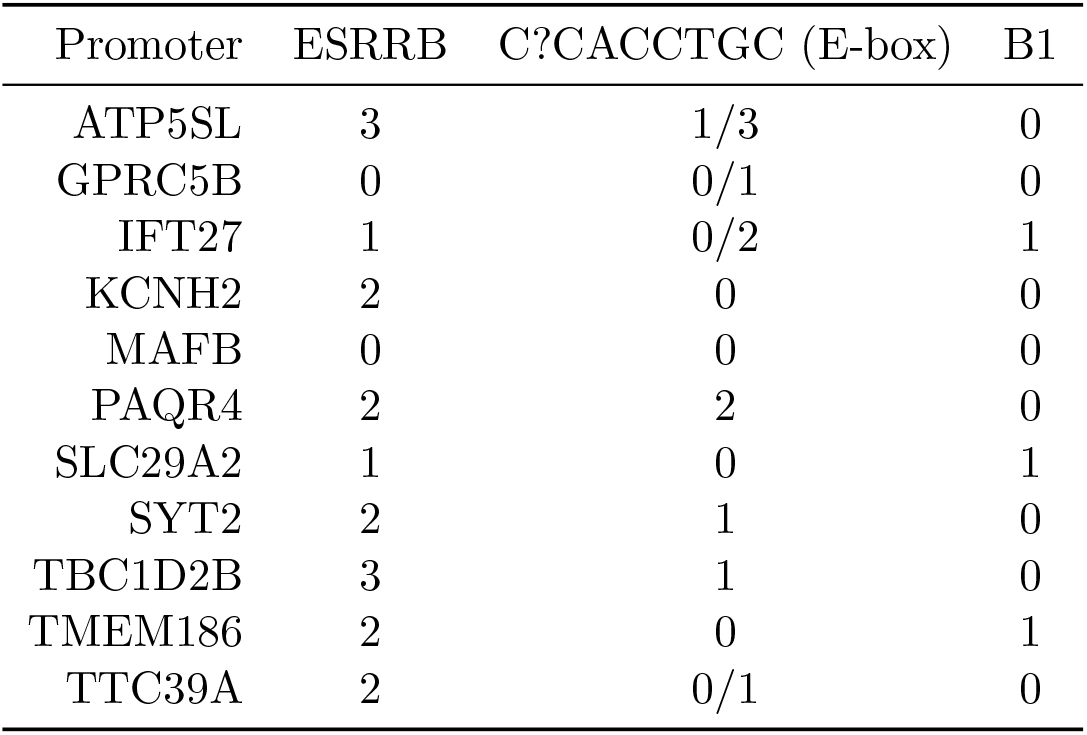
Selected candidate promoter regions contain different combinations of top motifs. Counts for the JASPAR motif ESRRB—the best JASPAR match to the top SArKS motif TGACCTTG— were assessed using fimo, while counts of the E-box sequence CCACCTGC or its reverse complement were assessed using simple string matching. If a promoter had additional matches to the substring CACCTGC (on either strand) omitting the initial C, a second count for this reduced match is indicated after a forward slash. Matches for the B1 SINE sequence were counted using `blast` requiring a minimum 90% sequence identity and 70 bp alignment length.

| Promoter | ESRRB | C?CACCTGC (E-box) | B1 |
| --- | --- | --- | --- |
| ATP5SL | 3 | 1/3 | 0 |
| GPRC5B | 0 | 0/1 | 0 |
| IFT27 | 1 | 0/2 | 1 |
| KCNH2 | 2 | 0 | 0 |
| MAFB | 0 | 0 | 0 |
| PAQR4 | 2 | 2 | 0 |
| SLC29A2 | 1 | 0 | 1 |
| SYT2 | 2 | 1 | 0 |
| TBC1D2B | 3 | 1 | 0 |
| TMEM186 | 2 | 0 | 1 |
| TTC39A | 2 | 0/1 | 0 |

### S3.3 Computational complexity and scalability of SArKS

One of the major motivations behind SArKS’ method of discovering motifs—searching for blocks of lexicographically similar suffixes derived predominantly from high-scoring sequences—lies in the scalability of suffix-based methods. The number of suffixes of a string (or set of strings) scales linearly with the length of the string(s) involved: as a result, the steps involved in the SArKS algorithm for identifying significant peaks scale linearly in both runtime and memory space with the combined size of the set of input sequences. We discuss this in more detail below. We then discuss the complexity of the later steps involved in extracting information regarding specific motif k-mers from the significant SArKS peak set.

The existing implementation of SArKS generates and then stores in memory the full suffix array of the concatenated sequence *x* = *ω*_0_ * …*ω*_n-1_: this step is asymptotically linear in the length of the concatenated sequence both in terms of runtime and memory (Kärkkäinen and Sanders, 2003). There is one caveat regarding the memory requirement here: the suffix array for a sequence of length l contains a permutation of the first l integers; while the length of this array is linear in l, the number of digits required to specify each integer grows logarithmically with l as well. An uncompressed suffix array (as used here) thus technically requires memory specified in bits scaling with llogl. Assuming the default use of 64-bit integers (as is done in the numpy-based python implementation we have used), however, memory will scale linearly for sequences of length up to ~ 1018 characters, far beyond current practical limits.

Given the inverted suffix array is yielding the value of the suffix array index *i* corresponding to the suffix array value si, the block array (Equation (3)) can be constructed in linear time and space (again in terms of the length of the concatenated sequence x) by (1) looping through the positions *s* in the concatenated string *x*, (2) checking whether the active block *b* needs to be incremented according to whether *s* ≥ *l_b+1_* (Section 2.1), and (3) filling in the position *i_s_* of the block array with the active block value *b*.

Kernel smoothing using a uniform kernel may be implemented in linear time by computing differences of cumulative sums (Equation (6)). The array of Gini impurity values (Equation (S4)) can be computed in linear time by successively computing the difference in consecutive values resulting from shifting the smoothing window by one position and updating the associated block frequencies Equation (S3). Identification of peaks (by comparing the score of each position to the scores of the two spatially adjacent positions) in the array of smoothed suffix scores *ŷi*, along with the filtering of the resulting peak set based on score threshold *θ* and Gini impurity threshold min, again requires time linear in the length of the concatenated sequence *x*. Similar remarks hold for the analogous spatial smoothing operations.

Permutation testing requires repetition of the above steps *R* times, where *R* is the number of permutations, and is hence still asymptotically linear in the length of the concatenated sequence *x*. While in principle parallelizable, each permutation will require its own smoothed (and, if desired, spatially smoothed) score array, so that parallelization requires memory linear in *R* * *|x|*.

Motif length selection according to Equation (7) could be naively implemented in *0*(*k_max_* k*) time per peak by directly comparing each suffix in the smoothing window to the suffix corresponding to the suffix array index around which the window is centered. In fact it is generally faster to use the suffix array to compute the suffix array index bounds for which the k-mer prefixing the central suffix is conserved (this may be done quite efficiently using the Burrows-Wheeler transform (Ferragina and Manzini, 2000); in our implementation of SArKS we have generally avoided this in order to reduce the memory requirements of the algorithm, favoring instead a slightly less efficient binary search approach). Either way, motif length selection generally requires time linear in the size of the peak set; in practice, when the peak set is large, this step can be relatively time consuming.

Merging of spatially adjacent k-mers originating within the same spatial smoothing window (Equation (S15)) may be computed in time linear in the size of the peak set times the length of the spatial smoothing window λ. In the case of large peak sets, this step can be time consuming as well.

## S4 Future directions

### S4.1 Gapped motif detection

While lexical sorting of suffixes assembles occurrences of the same k-mer into a block of adjacent index positions i, gapped motifs such as

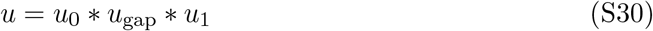

in which there is significant variability in the characters appearing within the internal substring *u_gap_* will be scattered into distinct subblocks dispersed within the larger superblock corresponding to their common prefix u_0_. This dispersion can dilute the apparent correlation *ŷi* between motif and score by mixing non-matching suffixes in with those corresponding to *u* within the range of the smoothing kernel.

While the technique described in Section S2.4 ameliorates this problem, it does not specifically focus on the important situation where a head motif u_0_ is always followed by the same tail motif *u1* after the variable region *u_gap_*. Such gapped motifs might be discovered using SArKS by first applying a relatively relaxed threshold *θ* (which may on its own admit many false positives) and then examining the tail sequences *u_gap_* * *u1* * • • • following it for evidence of an enriched sequence u_1_, removing candidate head sequences for which no such corresponding tails can be found. In this way, the ability of SArKS to detect motifs with particularly variable internal positions may be improved.

### S4.2 Other applications of SArKS

While we have tested SArKS as a method for identifying candidate cell type-specific regulatory motifs, it could also be applied to sequence motifs associated with state dependent changes in activated neurons of a single class as well as to differential gene expression in cancer and in specimens that have been exposed to varying physical or chemical stimuli. We also anticipate uses far afield from analysis of biological sequences, including motif discovery in time series data (Fu, 2011), or, by considering node or edge sequences produced by random walks, analysis of complex network structure (Masoudi-Nejad *et al*, 2012).

